# HIV-1 uncoating by release of viral cDNA from capsid-like structures in the nucleus of infected cells

**DOI:** 10.1101/2020.11.13.380030

**Authors:** Thorsten G. Müller, Vojtech Zila, Kyra Peters, Sandra Schifferdecker, Mia Stanic, Bojana Lucic, Vibor Laketa, Marina Lusic, Barbara Müller, Hans-Georg Kräusslich

## Abstract

HIV-1 replication commences inside the cone-shaped viral capsid, but timing, localization and mechanism of uncoating are under debate. We adapted a strategy to visualize individual reverse-transcribed HIV-1 cDNA molecules and their association with viral and cellular proteins using fluorescence and correlative-light-and-electron-microscopy (CLEM). We specifically detected HIV-1 cDNA inside nuclei, but not in the cytoplasm. Nuclear cDNA initially co-localized with a fluorescent integrase fusion (IN-FP) and the viral CA (capsid) protein, but cDNA-punctae separated from IN-FP/CA over time. This phenotype was conserved in primary HIV-1 target cells, with nuclear HIV-1 complexes exhibiting strong CA-signals in all cell types. CLEM revealed cone-shaped HIV-1 capsid-like structures and apparently broken capsid-remnants at the position of IN-FP signals and elongated chromatin-like structures in the position of viral cDNA punctae lacking IN-FP. Our data argue for nuclear uncoating by physical disruption rather than cooperative disassembly of the CA-lattice, followed by physical separation from the pre-integration complex.

## Introduction

Retroviral replication involves reverse transcription of the viral RNA genome and requires nuclear entry of the subviral complex to allow for chromosomal integration of the viral cDNA mediated by the viral integrase (IN) (Lusic & Siliciano, 2017). Whereas simple retroviruses require nuclear envelope breakdown during mitosis for productive replication, HIV-1 and other lentiviruses infect non-dividing cells, implying that the subviral complex can pass through the intact nuclear envelope (Suzuki & Craigie, 2007). Reverse transcription mediated by the viral reverse transcriptase (RT) is initiated in the cytoplasm, but recent evidence indicates that cDNA synthesis is completed inside the nucleus (Burdick et al., 2020; Dharan et al., 2020; Selyutina et al., 2020), at least in the case of HIV-1. The cytoplasm is a hostile environment for retroviral genome replication: exposure of cytoplasmic DNA to cellular nucleic acid sensors would lead to induction of innate immunity (Doitsh et al., 2014; Monroe et al., 2014), thereby aborting viral infection. The viral cone-shaped capsid apparently plays a central role in guiding and shielding (Rasaiyaah et al., 2013) the genome through the cytosolic environment (Campbell & Hope, 2015; Novikova et al., 2019). It consists of ~1,200-1,500 CA molecules assembled into hexamers and pentamers (Briggs et al., 2003), which have been shown to interact with components of the nuclear pore complex (NPC) (Dharan et al., 2016; Fernandez et al., 2019; Lee et al., 2010; Schaller et al., 2011; Zhou et al., 2011), implying a role for the CA-lattice in nuclear entry. However, the HIV-1 capsid, measuring ~60 nm at its wide end (Mattei et al., 2016), is considered to exceed the dimensions of the NPC channel with a reported maximal diameter of ~40 nm (von Appen et al., 2015). This implies that capsid uncoating should occur-at least partially - prior to nuclear entry, and various publications reported uncoating either in the cytoplasm (Cosnefroy et al., 2016; Mamede et al., 2017; Xu et al., 2013) or at the nuclear pore (Francis & Melikyan, 2018), with some evidence for cell type dependent differences. On the other hand, nuclear HIV-1 pre-integration complexes (PIC) were found to retain varying amounts of CA molecules (Bejarano et al., 2019; Burdick et al., 2020; Chin et al., 2015; Hulme et al., 2015; Stultz et al., 2017; Zila et al., 2019), at least in certain cell types (Zila et al., 2019), and recent reports indicating the presence of intact capsid lattice (Dharan et al., 2020; Selyutina et al., 2020) and capsid-like structures (Zila et al., 2020) inside the nucleus challenged the current models of early HIV-1 replication. Accordingly, the timing, subcellular localization, trigger and mechanism of HIV-1 capsid uncoating are still under debate.

Studying early HIV-1 replication is hampered by the fact that most cytoplasmic entry events appear to be non-productive in tissue culture (Klasse, 2015; Sanjuán, 2018). Therefore, characterization of individual subviral complexes containing viral cDNA with respect to their content, subcellular distribution and trafficking is required to shed light on the pathway of productive replication. Viral cDNA can be visualized in fixed cells using fluorescence in-situ hybridization (FISH) (Marini et al., 2015) or its derivatives using branched probes (Chin et al., 2015), but the harsh assay conditions destroy the native cellular environment and impair immunofluorescence analysis. Incorporation of the modified nucleoside 5-ethynyl-2’-deoxyuridine (EdU) allowed the detection of actively transcribing HIV-1 complexes by visualizing *de novo* synthesized viral DNA *via* click chemistry (Peng et al., 2015; Stultz et al., 2017), but this approach is also limited to fixed cells and cellular extraction precludes high resolution analysis (Müller et al., 2019). To overcome these limitations, we adapted a live cell compatible genetically encoded system (ANCHOR) that allows single molecule gene labeling (Germier et al., 2017; Saad et al., 2014), and has previously been applied for visualization of viral DNA (Blanco-Rodriguez et al., 2020; Komatsu et al., 2018; Mariamé et al., 2018). This system is based on the prokaryotic chromosomal partitioning system ParB-parS, where ParB (designated OR) specifically binds the parS seed sequence (designated ANCH). Multiple copies of parS introduced into the HIV-1 genome act as nucleation sites to oligomerize the fluorescently labelled OR protein when the reverse transcribed ANCH cDNA sequence becomes accessible to the fusion protein.

Here, we show that HIV-1 cDNA containing subviral complexes associated with CA are detected in the nucleus of infected cells, including primary CD4^+^ T cells. Over time, these nuclear complexes segregate from their CA content and the bulk of viral replication proteins, confirming nuclear uncoating. Using CLEM, we detected capsid-like structures at the position of nuclear IN-FP-containing complexes, whereas elongated, chromatin-like densities were observed at the position of viral cDNA punctae. Importantly, strong CA signals were observed on nuclear HIV-1 complexes in all cell types analyzed, indicating that prior failure to detect nuclear CA was largely due to masked epitopes.

## Results

### The ANCHOR system enables visualization of integrated and unintegrated HIV cDNA in the nucleus of infected cells

To test for retention of the ANCH sequence and efficiency of visualizing HIV-1 cDNA following reverse transcription, we stably transduced HeLa-based TZM-bl cells with different amounts of an ANCH containing HIV-1-based vector. These cell populations were subsequently transduced or transfected with an expression vector for eGFP.OR3 (Figure 1 - figure supplement 1a). Confocal microscopy revealed distinct eGFP.OR3 punctae in the nuclei of > 90 % of transfected cells (Figure 1 - figure supplement 1b-d). At low multiplicities of transduction with the ANCH-vector, where the majority of cells is expected to originate from a single integration event, we observed an average of 1.6 ± 0.29 and 1.4 ± 0.34 eGFP.OR3 punctae per nucleus (Figure 1 - figure supplement 1d). The number of punctae correlated with the multiplicity of transduction over a wide range (Figure 1 - figure supplement 1d). Of note, the eGFP.OR3 signal was stable for more than four weeks (when unintegrated viral cDNA species are degraded) suggesting that integrated viral DNA can be detected. Thus, the ANCH sequence is retained during reverse transcription, and this approach can be used to detect HIV-1 cDNA during the early replication phase.

Next, we introduced the ~1,000 bp ANCH sequence into the HIV-1 proviral plasmid pNLC4-3 (Bohne & Kräusslich, 2004) (HIV ANCH) replacing part of the *env* gene (Figure 1a). Virus-like particles were pseudotyped with the vesicular stomatitis virus G protein (VSV-G) or HIV-1 Env as indicated and also incorporated exogenously expressed IN (Albanese et al., 2008) tagged with a fluorescent marker (IN-FP) for visualization of subviral replication complexes. These particles were used to infect polyclonal TZM-bl cell populations stably transduced to express OR3 fused with either eGFP, mScarlet (Bindels et al., 2016) or the stainable SNAP-tag (Keppler et al., 2003); these cells also stably expressed fluorescently tagged Lamin B1 (LMNB1) to clearly distinguish nuclear and cytoplasmic events. Figure 1b shows TZM-bl eBFP2.LMNB1 and eGFP.OR3 expressing cells infected with HIV ANCH. Distinct infection-induced eGFP punctae were clearly detected in the nuclei of these cells, but were not observed in the cytoplasm where eGFP.OR3 was diffusely distributed (Figure 1b). Distinct nuclear eGFP punctae were not detected in uninfected cells.

**Figure 1.**
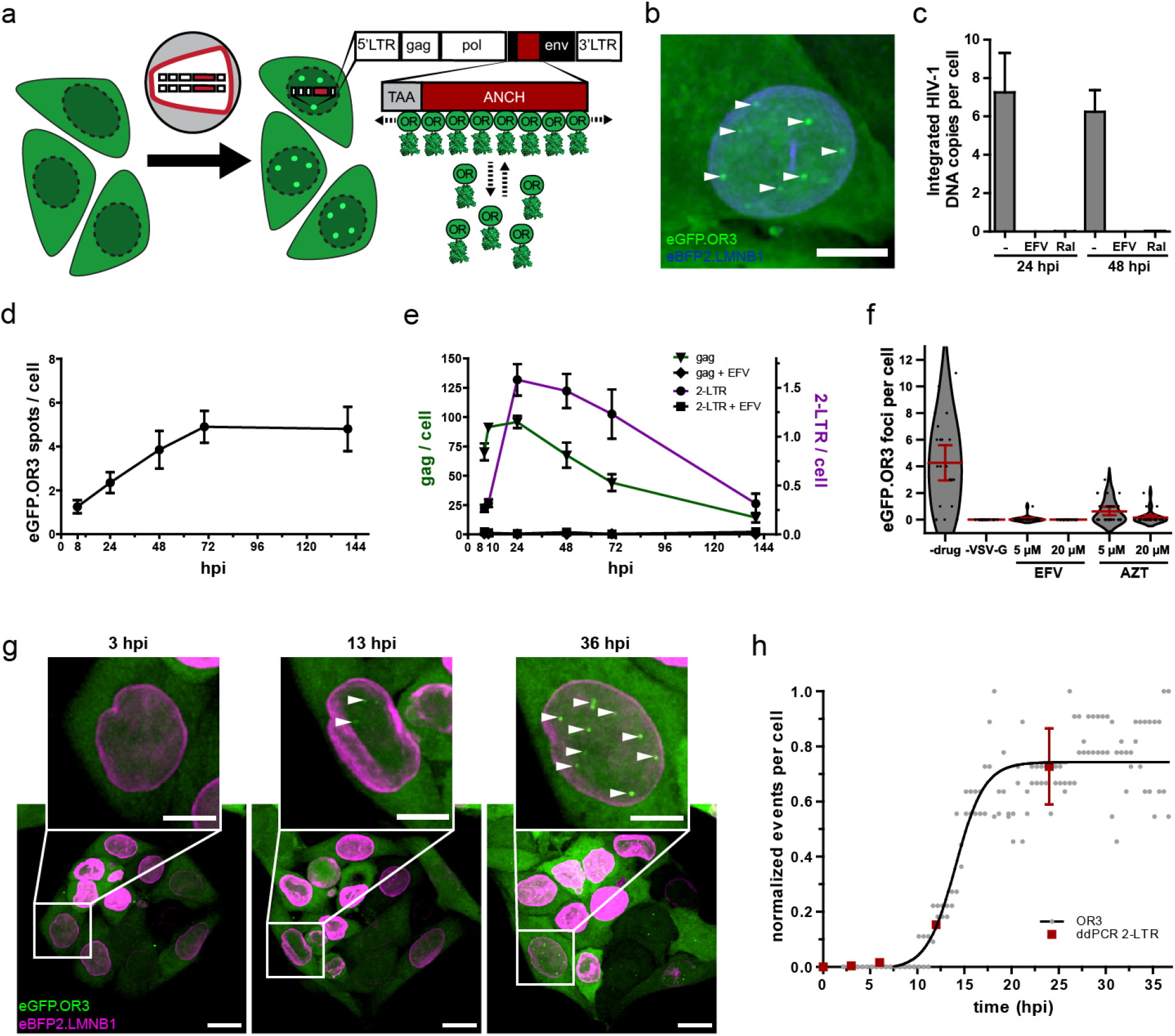
Visualization of HIV-1 dsDNA within the nucleus of infected cells. (**a**) Scheme of the ANCHOR dsDNA visualization system. Fluorescently tagged OR3 binds to the ANCH sequence on the viral dsDNA. (**b**) eGFP.OR3 punctae detected in the nuclei of infected cells. TZM-bl eBFP2.LMNB1 eGFP.OR3 cells were infected with VSV-G pseudotyped HIV-1_NL4-3_ ANCH (30 μUnits RT/cell, MOI 6), fixed at 55 h p.i. and imaged using SDCM. Six independent experiments with two independent virus preparations were performed. Maximum intensity projection (MIP) of a representative cell is shown. Note that some cells show one or two cytoplasmic, mostly perinuclear, infection-independent accumulations of the OR3 fusion protein, which we identified as multivesicular bodies (MVB) by CLEM (Figure 1 - figure supplement 3). Scale bar: 5 μm. (**c**) Quantification of integrated HIV-1_NL4-3_ ANCH provirus using nested Alu-LTR PCR. SupT1 cells were infected using VSV-G pseudotyped HIV ANCH (10 μU RT/cell). Raltegravir (RAL) or Efavirenz (EFV) were added at the time of infection as indicated. Data are shown for one experiment performed in biological triplicates; error bars represent SD. (**d,e**) eGFP.OR3 punctae are stable for more than 6 days while total and unintegrated DNA declined. Quantification of eGFP.OR3 punctae by SDCM (**d**), and *gag* and 2-LTR cDNA by ddPCR (**e**) over time within the same experiment. Infection was performed as in (**b**). (**d**) Data from one of two independent experiments (n = 20 cells per time point, error bars represent SEM). (**e**) Data from one experiment performed in biological triplicates (error bars represent SEM). (**f**) Detection of eGFP.OR3 punctae is HIV-1 cell fusion and reverse transcription dependent. TZM-bl eBFP2.LMNB1 and eGFP.OR3 cells were infected with VSV-G pseudotyped NNHIV ANCH (10 μU RT/cell) and imaged under live conditions at 27 h p.i‥ Control experiments were performed adding HIV-1 RT inhibitors EFV or azidothymidine (AZT) at time of infection or using particles lacking a fusion protein (“-VSV-G”). n = 20-29 cells were analyzed per sample and error bars represent 95 % CI; The graph shows data from one of three independent experiments. (**g,h**) Live cell imaging of vDNA punctae formation. TZM-bl eBFP2.LMNB1 and eGFP.OR3 cells were infected with NNHIV ANCH (30 μU RT/cell) and image acquisition by SDCM was initiated at 2 h p.i‥ 3D stacks were acquired every 30 min for 36 h. Representative data from one of four independent experiments are shown. MIP of a representative cell is shown. Scale bars: 10 μm (overview), 5 μm (enlargement). See Video 1. (**h**) Quantification of eGFP.OR3 punctae formation in cells from video shown in (g). Analyzed was the eGFP.OR3 punctae formation from two cells within the field of view (FOV) with each point representing the normalized amount of OR3 punctae per nucleus at the respective timepoint. Data were fit to a logistic growth model giving t_1/2_ = 14.1 ± 0.5 min. Detection of 2-LTR circles (mean and SEM) by ddPCR in biological triplicates is shown for TZM-bl cells infected with NNHIV WT.

To determine whether HIV ANCH cDNA became chromosomally integrated, we infected the human SupT1 T-cell line with HIV ANCH and analyzed the copy number of integrated proviral genomes by semi-quantitative Alu-PCR; this experiment could not be performed in TZM-bl cells, since these cells carry multiple HIV-1 LTR copies from prior lentiviral vector transductions. Integrated proviral DNA was readily detected in HIV ANCH infected SupT1 cells, but was not observed when infection was performed in the presence of an RT-or IN-inhibitor (Figure 1c). Similar to TZM-bl cells, SupT1 cells also showed nuclear eGFP.OR3 punctae following infection with HIV ANCH (see below, Figure 6 - figure supplement 1a). We then determined the number of eGFP.OR3 punctae in TZM-bl cells using confocal microscopy of cells fixed at different time points after infection with HIV ANCH (Figure 1d) and performed parallel quantitation of total HIV-1 cDNA and 2-LTR circles (representing unintegrated nuclear HIV-1 cDNA) using digital droplet PCR (ddPCR) (Figure 1e). Total cDNA levels became saturated at 10 hours post infection (h p.i.) and 2-LTR circles peaked at 24 h p.i.; both species strongly declined over the following five days (Figure 1e). eGFP punctae, on the other hand, increased over the first 72 h, but then remained stable over the following three days despite the observed loss of HIV-1 cDNA species (Figure 1d). Taken together, these results clearly indicated that integrated HIV-1 DNA can be detected using the ANCHOR system. To determine whether detection of integrated proviral copies may be influenced by RNA transcription at the respective site, we compared the number of eGFP.OR3 punctae in HIV ANCH infected TZM-bl cells treated with the CDK9/p-TEFb inhibitor Flavopiridol or solvent. No difference was observed (Figure 1 - figure supplement 2a-c), suggesting that the dynamic nature of OR3 recruitment does not interfere with transcription.

Next, we generated a non-infectious derivative of HIV ANCH termed NNHIV ANCH to allow live cell imaging outside the BSL3 facility. NNHIV ANCH is based on the previously reported plasmid NNHIV that carries mutations in the active site of IN and a deletion in the *tat* gene (IN_D64N/D116N_ tat_Δ33-64bp_), thereby retaining reverse transcription and nuclear import, but lacking the capacity to integrate and transcribe (Zila et al., 2020). TZM-bl eGFP.OR3 cells infected with VSV-G pseudotyped NNHIV ANCH also showed nuclear eGFP.OR3 punctae (Figure 1f-h) indicating that unintegrated HIV-1 cDNA is detected by the ANCHOR system as well. eGFP punctae were not detected when cells were treated with NNHIV ANCH lacking VSV-G and were lacking or strongly reduced in the presence of the RT inhibitors efavirenz (EFV) or azidothymidine (AZT) (Figure 1f).

To investigate the dynamics of appearance of eGFP.OR3 punctae in NNHIV ANCH infected TZM-bl cells, we performed live cell imaging experiments using spinning disk confocal microscopy (SDCM). The onset of marker recruitment to viral cDNA in the nucleus was observed at 7-8 h p.i., while the half-maximal signal was reached between 13 and 15 h p.i. (Figure 1g, h; Video 1). Again, no infection-induced eGFP.OR3 punctae were detected in the cytosol of infected cells. The onset of nuclear HIV-1 2-LTR detection using ddPCR coincided with the appearance of eGFP.OR3 punctae (Figure 1h). Formation of both 2-LTR circles and eGFP.OR3 punctae requires viral cDNA to be synthesized and accessible to proteins not present in the subviral replication complex (2-LTR formation requires nuclear NHEJ components and ligase IV (Li et al., 2001)). Accordingly, the lack of cytoplasmic eGFP.OR3 punctae may be due to incomplete cDNA synthesis in the cytoplasm and/or to cDNA only becoming accessible to the fusion protein upon full capsid uncoating in the nucleus. To address this question we focused on the timing and quantification of reverse transcription in the described system.

### Nuclear eGFP.OR3 punctae contain higher amounts of *de novo* synthesized HIV-1 cDNA than nuclear IN-positive complexes not recruiting eGFP.OR3

FISH analysis revealed a large amount of RT products in the cytoplasm of infected cells, where no eGFP.OR3 punctae were observed (Figure 2a). When infecting TZM-bl cells with equal amounts of NNHIV ANCH for 24 h, we observed ~130 *gag* (Figure 2b) and two to three 2-LTR cDNA molecules (Figure 2c) per cell and ~4 nuclear eGFP.OR3 punctae per cell (Figure 1f). These results clearly showed that the majority of late RT products - including all cytoplasmic products - are not associated with eGFP.OR3.

**Figure 2.**
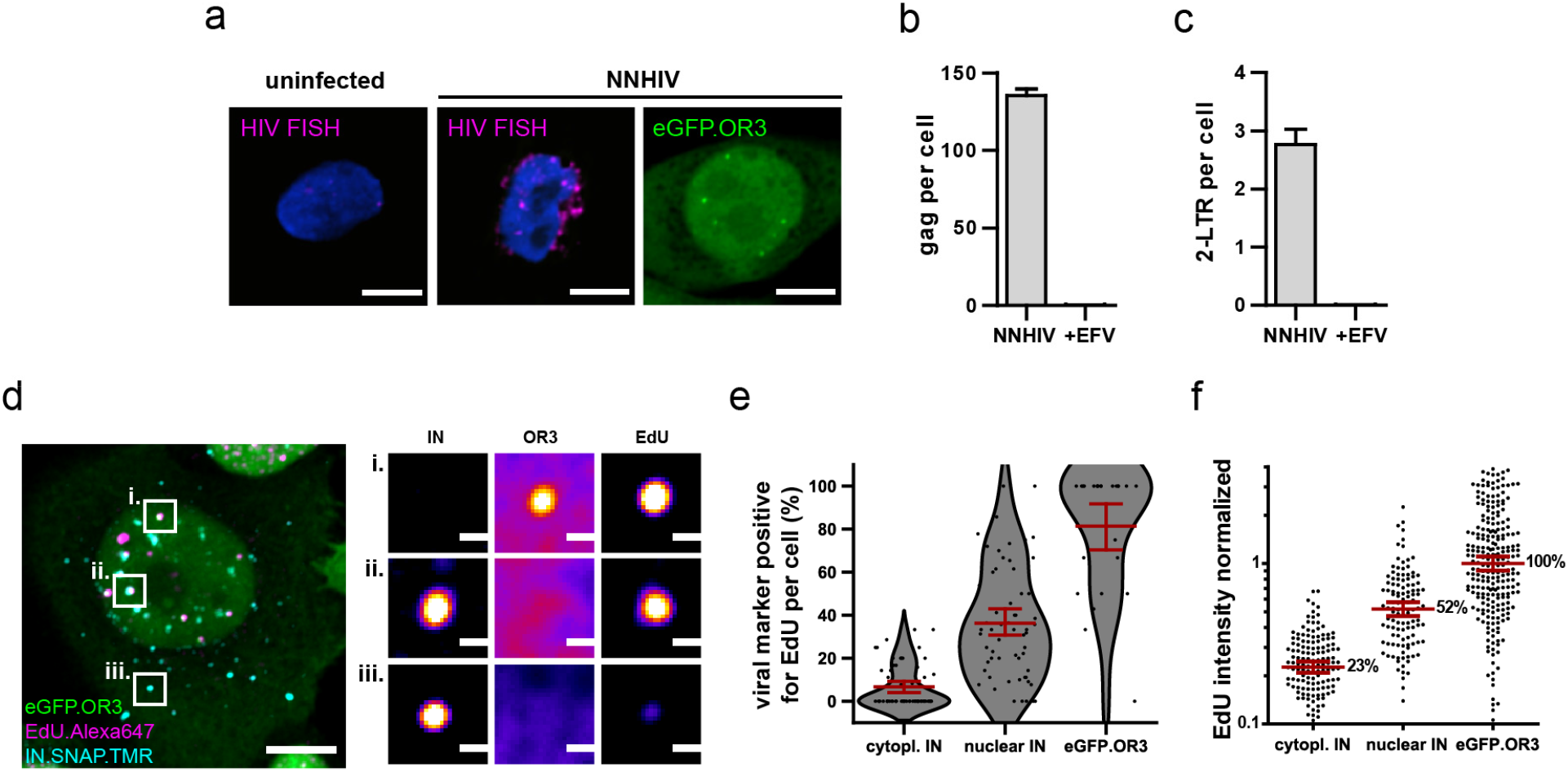
Viral cDNA in cytosolic complexes does not recruit OR3 proteins and contains less DNA compared to nuclear complexes. (**a**) HIV FISH staining of TZM-bl cells infected with VSV-G pseudotyped NNHIV ANCH (30 μU/RT cell; 24 h p.i.). A cell showing nuclear eGFP.OR3 signals from a different experiment is shown for comparison. Scale bars: 5 μm. (**b,c**) ddPCR quantification of late RT products (*gag* region; **b**) and 2-LTR circles (**c**). TZM-bl cells were infected using the same amount of VSV-G pseudotyped NNHIV ANCH (10 μU RT/cell, 24 h p.i.) as in Figure 1f. Mean and SEM of one experiment performed in biological triplicates are shown. (**d**) TZM-bl eBFP2.LMNB1 and eGFP.OR3 expressing cells were infected with VSV-G pseudotyped and IN.SNAP.TMR labelled NNHIV ANCH (30 μU RT/cell) in the presence of EdU. Cellular DNA synthesis was blocked by aphidicolin (APC). At 24 h p.i. cells were fixed and click labelled. Two independent experiments were performed in biological triplicates. A 2 μm MIP of a representative cell and three enlarged single z slices are shown. Scale bars: 5 μm (MIP) and 1 μm (enlargements). (**e,f**) Quantification of data from the experiment described in (**d**). Pooled data from one experiment performed in biological triplicates are shown, with data points representing individual cells (**e**) or subviral complexes (**f**); error bars represent 95 % CI. (**e**) Percentage of EdU positive viral marker spots (IN or OR3) per cell. (**f**) Intensity of EdU signals associated with the respective viral marker. Data were normalized to the mean signal of eGFP.OR3 punctae. Differences are statistically significant (p < 0.0001; two-tailed Student’s t-test).

Progress of reverse transcription was also assessed at the single particle level. For this, we infected cells with NNHIV ANCH carrying fluorescently tagged IN-FP in the presence of EdU followed by fluorescent click labelling of newly synthesized DNA at different time points p.i‥ Cellular DNA synthesis was inhibited by the DNA polymerase α/δ inhibitor aphidicolin (APC). Co-localization with EdU was observed for 7 % of IN-FP positive structures in the cytoplasm (95 % CI of mean: 4-9 %) and for 36 % of IN-FP positive complexes in the nucleus (95 % CI of mean: 29-43 %). Importantly, 81 % of nuclear eGFP.OR3 punctae were positive for EdU (95 % CI of mean: 70-93 %; Figure 2d, e). While some nuclear EdU-positive complexes were positive for both the IN-FP and eGFP.OR3, we also observed HIV-1 cDNA containing complexes that were only positive for either IN-FP or eGFP.OR3 (Figure 2d, panels i and ii). The average EdU signal was lower on cytoplasmic than on nuclear HIV-1 complexes and was highest on eGFP.OR3 punctae. When setting the EdU signal for eGFP.OR3 punctae to 100 %, the relative EdU signal was significantly reduced to 23 % (p < 0.0001) on cytoplasmic IN-positive complexes and to 52 % (p < 0.0001) on nuclear IN-positive complexes (Figure 2f). These observations support recent reports that reverse transcription is completed inside the nucleus (Burdick et al., 2020; Dharan et al., 2020; Selyutina et al., 2020) and indicate that the viral cDNA only becomes detectable to eGFP.OR3 when reverse transcription has been completed.

### HIV cDNA separates from IN-fusion proteins in the nucleus of infected cells

The low degree of co-localization between the fluorescent IN fusion protein and eGFP.OR3 on EdU231 positive nuclear punctae at 24 h p.i. (Figure 2d) prompted us to analyze the relative distribution of both fluorescent proteins in a time resolved manner after NNHIV ANCH infection. At 8 h p.i., 70 ± 11 % of nuclear eGFP.OR3 punctae were also positive for IN.SNAP (Figure 3a-c). This co-localization was largely lost at 24 h p.i. with only 14 ± 6 % of nuclear eGFP.OR3 punctae positive for IN.SNAP (Figure 3a-c). Strikingly, IN.SNAP punctae were often observed in close vicinity of eGFP.OR3 punctae at this later time point (Figure 3a, right panel), suggesting that they may have separated from a common complex. Similar results were observed for HIV-1 ANCH (Figure 3 - figure supplement 1a), an integration competent lentiviral vector containing ANCH (Figure 3 - figure supplement 1b), and when particles were pseudotyped with HIV-1 Env instead of VSV-G (Figure 3 - figure supplement 1c). These results showed that the observed phenotype was not dependent on the cytosolic entry pathway or on integration competence. To directly address the possibility of separation of the proviral cDNA from IN.SNAP, we performed live cell imaging of infected cells. We observed gradual loss of the IN.SNAP signal correlating with increased eGFP.OR3 recruitment and eventual separation of eGFP.OR3 punctae and IN.SNAP containing complexes (Figure 3d, e, Video 244 2). Of note, we occasionally observed consecutive appearance of two individual eGFP.OR3 punctae and their subsequent separation from the same IN.SNAP complex (Video 3), suggesting that single diffraction limited IN.SNAP punctae may correspond to multiple cDNA containing subviral HIV-1 complexes.

**Figure 3.**
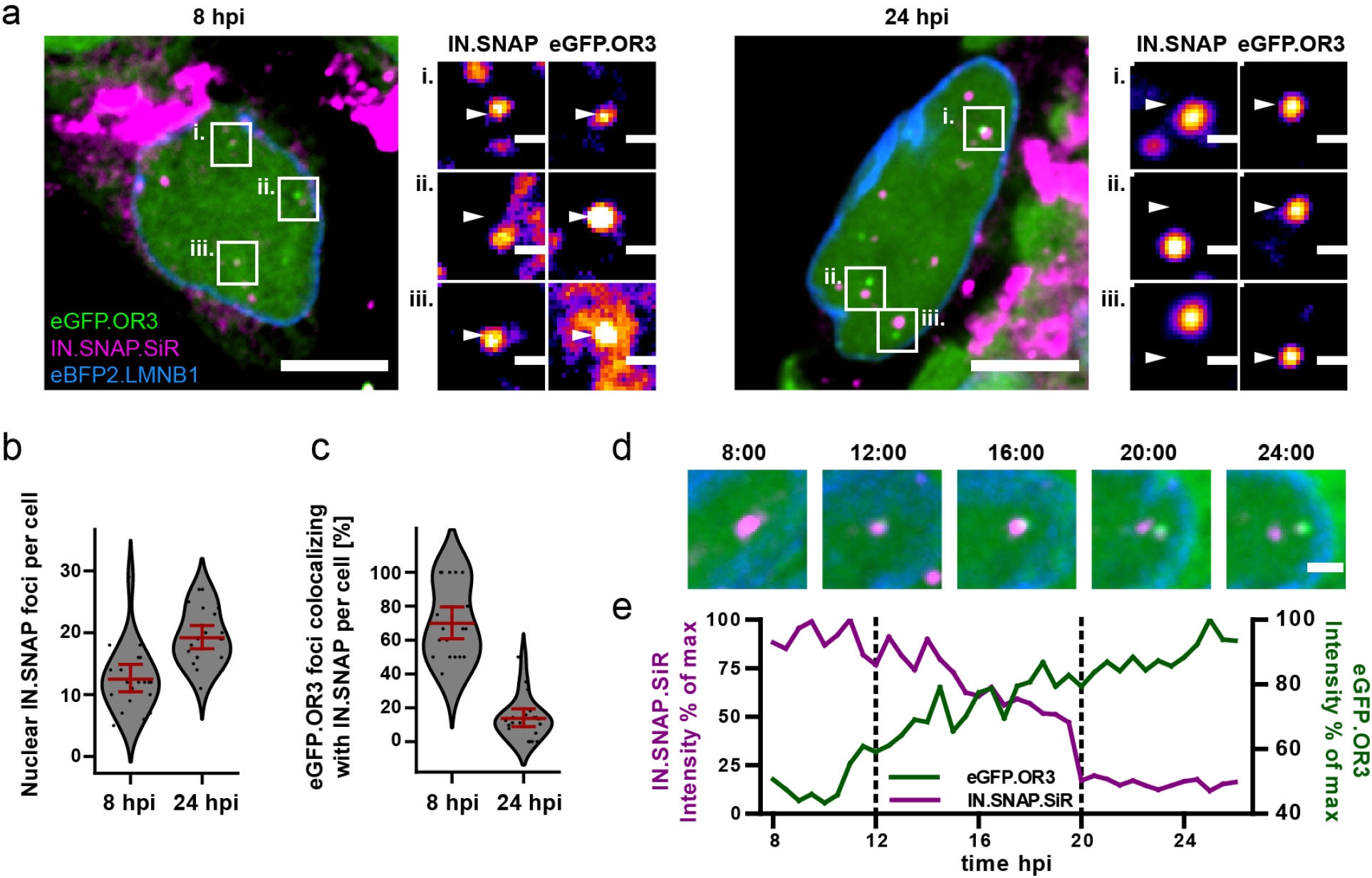
Separation of viral cDNA from IN marker within the nucleus. (**a**) TZM-bl eBFP2.LMNB1 eGFP.OR3 cells were infected with VSV-G pseudotyped and IN.SNAP.SiR labelled NNHIV ANCH particles (30 μU RT/cell), fixed at 8 and 24 h.p.i as indicated and imaged by SDCM. A 1 μm MIP of a representative cell and three enlarged single z slices are shown. Scale bars: 10 μm (MIP) and 1 μm (enlargements). The nuclear background of eGFP.OR3 was subtracted in enlargements for clarity. The figure shows representative images from one of three independent experiments. (**b**) Number of nuclear IN.SNAP punctae/cell detected at the indicated time p.i.; 13 ± 3 punctae/cell (8 h p.i.; ± 95 % CI) and 19 ± 2 punctae/cell (24 h p.i.; ± 95 % CI). Error bars represent 95 % CI. (**c**) eGFP.OR3 and IN.SNAP co-localization observed over time; 70 ± 11 % (8 h p.i.; ± 95 % CI) and 14 ± 6 % (24 h p.i.; ± 95 % CI). Error bars represent 95 % CI. (**d**) Live imaging of IN.SNAP.SiR (magenta) and eGFP.OR3 (green) signal separation within the nucleus. eBFP2.LMNB1 is shown in blue. The figure shows individual frames from Video 2 (4.5 μm MIP) recorded at the indicated times (h:min p.i.). Scale bar: 2 μm; one of three independent experiments. Also see Video 3. (**e**) Relative intensities of IN.SNAP.SiR and eGFP.OR3 fluorescence detected at the eGFP.OR3 focus in (**d**). The plot comprises data corresponding to the position of the respective IN.SNAP focus recorded before the appearance of eGFP.OR3 (8-11 h p.i.). The area between the dotted lines corresponds to the period of colocalization between the major parts of the IN.SNAP signal and the eGFP.OR3 signal.

### Nuclear IN punctae exhibit a strong signal for CA and CPSF6 and correspond to clustered subviral particles

The observation that the IN.SNAP signal remained as a distinct cluster after separation of the eGFP.OR3-associated viral cDNA suggested that these clusters constitute a stable complex potentially containing other viral and cellular proteins held together by a scaffold. The viral capsid or a capsid-derived structure would be an obvious candidate for such a scaffold, but earlier studies detected no or only weak CA IF signals on nuclear HIV-1 complexes in HeLa derived cell lines (Peng et al., 2015), while a recent study reported strong nuclear signals for CA fused to GFP (Burdick et al., 2020). We decided to revisit this issue, since we had previously observed that CA immunostaining efficiency in the nucleus strongly depended on fixation and extraction conditions (Zila et al., 2019). Following extraction with methanol we indeed consistently detected a clear CA-specific signal co-localizing with most IN-positive punctae inside the nucleus (Figure 4a, top panel). Furthermore, these IN-positive punctae were also strongly positive for the host protein cleavage and polyadenylation specificity factor 6 (CPSF6) that binds specifically to the hexameric CA lattice (Figure 4a, bottom panel). Cells in this experiment had been treated with APC to arrest the cell cycle and prevent entry into mitosis, indicating that HIV-1 capsids or capsid-derived structures are able to transit the nuclear pore in HeLa derived cells, as had been reported for terminally differentiated macrophages (Bejarano et al., 2018; Stultz et al., 2017). To further analyze the ultrastructure of these nuclear CA-containing complexes, we employed stimulated emission depletion (STED) nanoscopy and correlative light and electron microscopy (CLEM).

**Figure 4.**
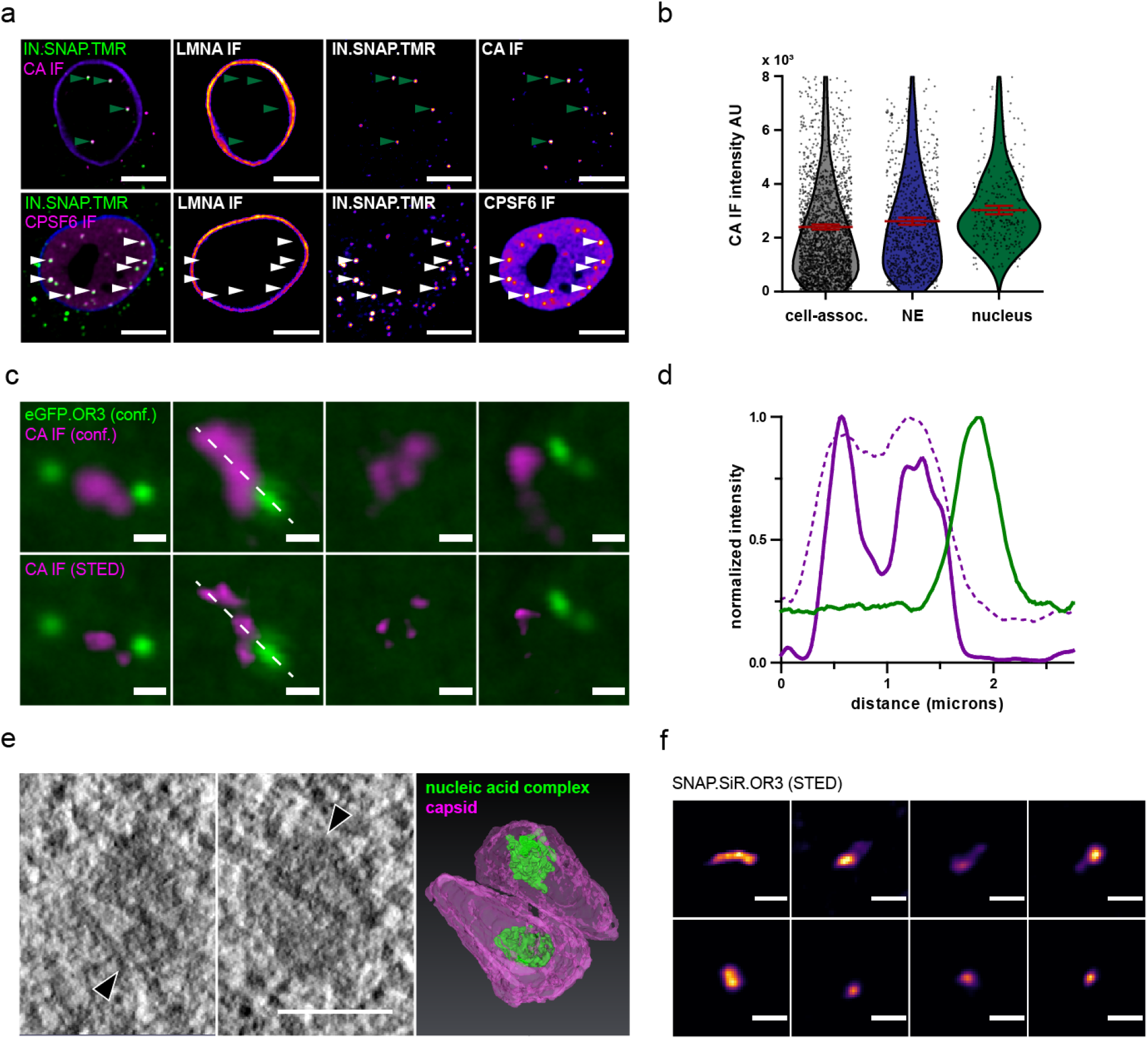
HIV-1 capsids cluster within the nucleus of HeLa derived cells. (**a**) IF detection of HIV-1 CA (top panel) and CPSF6 (bottom panel) in the nuclei of TZM-bl cells treated with APC and infected with VSV-G pseudotyped NNHIV ANCH labelled with IN.SNAP.TMR (30 μU RT/cell) and fixed at 24 h p.i‥ Data from one of three independent experiments are shown. Scale bars: 10 μm. (**b**) Quantification of CA intensities at IN.SNAP.TMR spots. Dots represent individual subviral complexes localized at the cytosol/plasma membrane (cell-assoc.), the NE or the nucleus; error bars represent SEM. Cells from five fields of view were analyzed (n = 2,610, 932 and 286 particles, respectively). (**c**) STED nanoscopy of CA accumulation at diffraction limited spots. Shown are four examples of nuclear CA and OR3 signals. TZM-bl eBFP2.LMNB1 and eGFP.OR3 cells were infected as in (**a**) without APC treatment; bottom panel shows STED microscopy of CA signals. Scale bars: 500 nm (**d**) Intensity profiles measured along the dashed white line in (**c**) normalized to the respective maximal value. Magenta, CA intensites in STED mode; dashed magenta, CA intensites in confocal mode; green, eGFP. OR3 intensities in confocal mode. (**e**) Electron tomography of NNHIV capsids detected in the nucleus. eGFP.OR3 expressing TZM-bl cells were infected with VSV-G pseudotyped NNHIV ANCH labelled with IN.SNAP.SiR (30 μU RT/cell) and high pressure frozen at 24 h p.i. Left and middle panels show slices through a tomographic reconstruction at the position within the nucleus correlated to an IN.SNAP spot (negative for eGFP.OR3). Arrowheads point to two closely associated cone-shaped structures where the wide end of one cone is oriented towards the narrow end of the other cone. Scale bar: 100 nm. The right panel shows the segmented and isosurface rendered structure of these cones. Magenta, capsid; green, nucleic acid containing replication complex. See Video 4. (**f**) STED nanoscopy of SNAP.OR3 expressing TZM-bl cells infected with VSV-G pseudotyped NNHIV ANCH (30 μU RT/cell) and fixed at 24 h p.i‥ Shown is a selection of eight nuclear SNAP.OR3 signals stained with O6-benzylguanine (BG)-SiR for 30 min prior fixation and analyzed by STED nanoscopy. Note that nuclear background of SNAP.OR3 has been subtracted for clarity. Scale bars: 300 nm.

The mature HIV-1 capsid contains ca. 50 % of the total CA content of the intact virion and post-fusion cytoplasmic capsids therefore exhibit a weaker CA signal than complete virions (Zila et al., 2019). Accordingly, CA specific immunofluorescence would be expected to be lower for nuclear complexes compared to cell-associated particles that represent a mixture of post-fusion particles and complete particles at the plasma membrane or endocytosed in the cytosolic region. However, the observed intensity of the CA signal on nuclear HIV-1 complexes was slightly higher than that observed for cell-associated particles (Figure 4b). This may be explained by exposure of additional epitopes due to capsid remodelling and/or by clustering of capsid-derived structures in a single diffraction limited spot. In order to investigate the latter possibility, we used STED nanoscopy to resolve individual subviral structures with a resolution of < 50 nm. Multiple individual CA signals in close vicinity to each other could be resolved within the area of a single focus detected in confocal mode (Figure 4c and d). For a more detailed analysis of the associated structures, we performed CLEM as described in the following section. IN.SNAP-positive and eGFP.OR3-negative nuclear punctae detected by fluorescence microscopy could be correlated with single or multiple electron-dense cone-shaped structures, whose shapes and dimensions closely resembled mature HIV-1 capsids (Figure 4e, Video 4). STED nanoscopy of nuclear SNAP.OR3-positive punctae corresponding to viral cDNA identified elongated structures (Figure 4f); in this case, only a single object was resolved by STED nanoscopy at each diffraction-limited position.

### Ultrastructure analysis reveals conical and elongated structures in the position of IN punctae and OR3 punctae, respectively

The findings described above suggest that apparently intact conical HIV-1 capsids can access the nucleoplasm, where reverse transcription is completed followed by separation of the viral cDNA from the bulk of viral proteins including CA. To determine the ultrastructure of the observed subviral complexes, we performed CLEM analysis. For this, we employed a TZM-bl mScarlet.OR3 cell line (Figure 3 - figure supplement 1d), because the mScarlet fluorescence signal was best retained upon plastic embedding. Cells were infected with VSV-G pseudotyped NNHIV ANCH carrying IN.SNAP.SiR, thus allowing direct high-pressure freezing of the sample without pre-fixation. Cells were vitrified at 24 h p.i. and thin sections were prepared after freeze-substitution and plastic embedding. Samples retained fluorescence for mScarlet.OR3 and IN.SNAP.SiR. Multi-channel fluorescent Tetraspeck markers were used for correlation and we identified positions corresponding to mScarlet.OR3 and IN.SNAP.SiR signals, respectively (Figure 5a-c). These positions were imaged using electron tomography (Figure 5d-i). At positions correlating to IN positive punctae (lacking mScarlet.OR3), we detected cone-shaped structures resembling HIV-1 capsids (Figure 4e) as well as less defined structures consistent with damaged cones or remnants of capsids (Figure 5g ii, h ii). Electron-dense material most likely corresponding to nucleic acid was visible inside most cone-shaped structures (Figure 4e and 5i top left panel, black arrowhead), whereas capsid remnant-like structures mostly lacked interior densities (Figure 5g ii. and Figure 5h ii.). In contrast, tomograms that correlated to positions of mScarlet.OR3 signals (lacking IN.SNAP.SiR signal) showed no defined conical or remnant-like structure. Instead, elongated densities were observed (Figure 5g i. and Figure 5h i.), in line with the findings from STED nanoscopy (see Figure 4f). These structures consisted of linked globular densities (with ~ 30 nm in diameter) resembling the appearance of chromatin, with lengths of ca. 100 nm (Figure 5h i.). One observed structure spanned 200-300 nm in length (Figure 5g i.).

**Figure 5.**
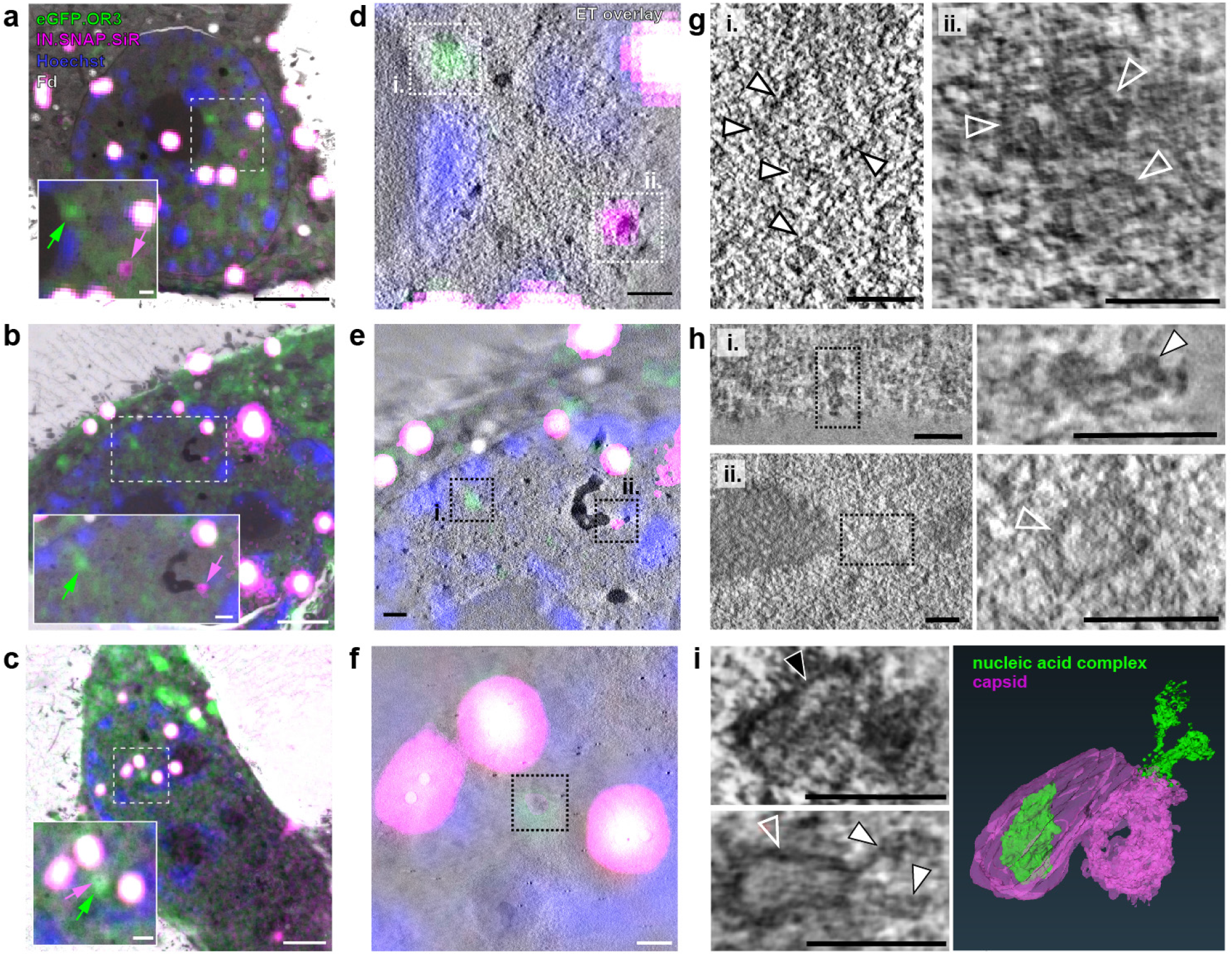
CLEM-ET analysis of IN and OR3 punctae inside the nucleus of infected HeLa derived cells. TZM-bl cells expressing mScarlet.OR3 were infected with VSV-G pseudotyped and IN.SNAP.SiR-labeled NNHIV ANCH (30 μUnits RT/cell). At 24 h p.i., cells were cryo-immobilized by high pressure freezing, freeze substituted and further processed for CLEM and ET. (**a-c**) CLEM overlays (with enlargements) of EM sections of cells expressing mScarlet.OR3 (green), infected with NNHIV ANCH IN.SNAP.SiR (magenta), post-stained with Hoechst (blue) and decorated with multi-fluorescent fiducials (Fd; white) for correlation. Positions of intranuclear spots positive for mScarlet.OR3 (green arrows) and IN.SNAP.SiR (magenta arrows) are indicated. Enlargements of area in dashed boxes is shown at the lower left of each panel. (**d-f**) CLEM-ET overlay of regions enlarged in (**a-c**). (**g-i**) Computational slices from tomographic reconstructions at the correlated positions boxed in (**d-f**). (**g**) Left, white arrowheads point to a filamentous structure corresponding to an mScarlet.OR3 (and IN.SNAP negative) spot boxed in (**d**; i.). Right, open white arrowheads indicate three capsid-reminiscent structures correlating to the IN.SNAP.SiR spot boxed in (**d**, ii.). (**h**) Top panels show a chromatin-like density, consisting of apparently linked globular structures, correlating to the mScarlet.OR3 positive and IN.SNAP.SiR negative spot boxed in (**e**; i.). Lower panels show the morphology of an empty open structure correlating to the IN.SNAP.SiR positive, mScarlet. OR3 negative spot boxed in (**e**; ii.). (**i**) Morphology of structures clustering at the position indicated by co-localizing mScarlet.OR3 and IN.SNAP.SiR spots boxed in (**f**). Top left, black arrowhead indicates an apparently intact capsid with density inside the cone. Bottom left, the open white arrowhead indicates an apparently empty cone-like structure. Note an elongated density (filled white arrowhead) protruding from the narrow end of the cone. The right panel shows the segmented and isosurface rendered structures shown on the left. See Video 5. Scale bars: 2.5 μm for overviews (**a-c**), 500 nm for enlargements (**a-c**), 250 nm (**d-f**), and 100 nm (**g-i**).

One complex visualized in these experiments correlated to both IN.SNAP.SiR and mScarlet.OR3. The corresponding tomogram revealed a dense cluster of three capsid-related structures (Figure 5i). One of these conical structures lacked interior density (Figure 5i, bottom left panel, open arrowhead) and appeared to be connected with an adjacent elongated density (filled white arrowheads) that appeared to protrude from the narrow end of the cone (Video 5, Figure 5i right panel). Taken together with the observations from live cell imaging, we speculate that this structure might represent a subviral complex captured in the process of or shortly after capsid uncoating.

### Nuclear CA and segregation of HIV-1 cDNA from the bulk of viral proteins are also observed in HIV-1 infected primary CD4^+^ T cells and monocyte-derived macrophages

In order to validate our findings in more relevant cell types, we adapted the system to the T cell line SupT1 and to primary CD4^+^ T cells and primary monocyte-derived macrophages (MDM). Of note, nuclear CA immunofluorescence signals have been detected previously in MDM (Bejarano et al., 2019), but not or only weakly in T cell lines (Zila et al., 2019) or primary T cells. Infection of an eGFP.OR3 expressing SupT1 cell line (Figure 6 - figure supplement 1) or of primary activated CD4^+^ T cells transduced to express eGFP.OR3 (Figure 6a) with NNHIV ANCH showed nuclear OR3 punctae and separation of IN-FP and OR3 punctae as observed for TZM-bl cells. In accordance with previous observations (Zila et al., 2019), no or weak CA signals were detected co-localizing with IN.SNAP foci in T cells (Figure 6a), even after applying methanol extraction. However, very strong signals for CPSF6, which binds to the hexameric CA lattice (Price et al., 2012), were detected at positions of the IN-FP (Figure 6b top panel). This observation led us to speculate that the underlying capsid lattice is masked by the dense coat of CPSF6 in these cells and is thus not accessible to antibody detection.

**Figure 6.**
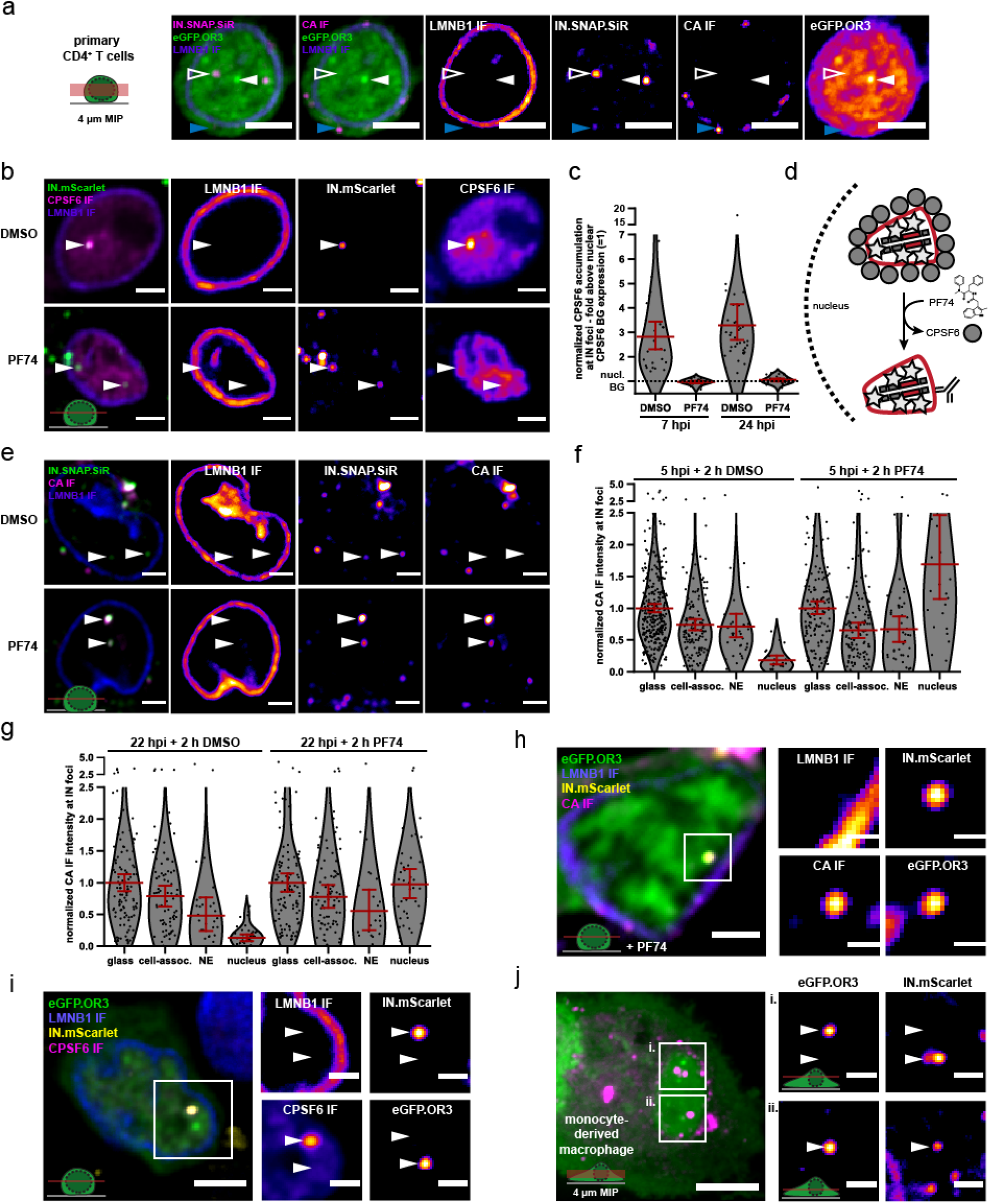
HIV DNA separates from IN/CA/CPSF6 positive structures in primary CD4^+^ T cells and MDM. (**a**) Activated CD4^+^ T cells were transduced with a lentiviral vector expressing eGFP.OR3. After 48 h, cells were infected using VSV-G pseudotyped NNHIV ANCH labelled with IN.SNAP.SiR (30 μU RT/cell) for 24 h before fixation and methanol extraction (as in Figure 4a-b). Arrowheads indicate the positions of a nuclear (left, open white) and cytoplasmic (bottom, blue) IN.SNAP particle as well as a nuclear eGFP.OR3 focus (right, filled white). A representative image of one of three independent experiments is shown. Scale bars: 5 μm. (**b-g**) Addition of PF74 (15 μM) after nuclear import enables immuno-detection of strong CA signals in CD4^+^ T cells. Activated primary CD4^+^ T cells were infected using VSV-G pseudotyped NNHIV ANCH IN.mScarlet or IN.SNAP.SiR (30 μU RT/cell). PF74 or DMSO was added after 5 or 22 h for another 2 h prior to fixation and methanol extraction. Shown is one of two independent experiments performed with cells from three different blood donors. (**b**) PF74 displaces CPSF6 from IN spots. Shown are single z slices from cells fixed at 24 h p.i. and immunostained for CPSF6. Scale bars: 3 μm (**c**) Quantification of CPSF6 intensity at IN spots. IN objects were segmented in 3D data sets. The CPSF6 mean intensity of these volumes was quantified as described in materials and methods. Dots represent single subviral complexes and error bars represent 95 % CI. (**d**) Scheme visualizing PF74-mediated displacement of CPSF6. (**e**) PF74 enables CA IF detection at IN spots. Shown are single z slices from cells fixed at 24 h p.i. and immunostained for HIV-1 CA. Scale bars: 3 μm. (**f,g**) Quantification of CA intensities at IN spots at 7 h (**f**) and 24 h (**g**) p.i‥ IN positive objects were segmented in 3D. The CA mean intensity of these volumes was quantified and normalized to the CA intensity of IN positive objects located on glass inside the same field of view. Error bars represent 95 % CI. P values of differences between DMSO and PF74 treatments (two-tailed Student’s t-test): (glass = 1.000 (**f**, not significant (ns)), 1.000 (**g**, ns); cell-assoc. = 0.2684 (**f**, ns), 0.9427 (**g**, ns); NE = 0.7884 (**f**, ns), 0.7514 (**g**, ns); nucleus = < 0.0001 (**f**, significant), < 0.0001 (**g**, significant). (**h**) CA positive structure colocalizing with IN and vDNA markers inside the nucleus of CD4^+^ T cells. PF74 was added for 2 h prior to fixation at 24 h p.i‥ Nuclear background of OR3 was subtracted in enlargements for clarity (in **h-j**). Shown is a single z slice, scale bars: 5 μm (overview) and 1 μm (enlargement). (**i**) CPSF6 remains colocalized with the IN signal. Shown is a single z slice, scale bars: 5 μm (overview) and 2 μm (enlargement). (**j**) MDM were transduced using a Vpx containing eGFP.OR3 expressing lentiviral vector. After 72 h cells were infected using VSV-G pseudotyped NNHIV IN.mScarlet (120 μU RT/cell), fixed and imaged at 24 h p.i‥ Shown is a 4 μm maximum intensity projection. Scale bar: 5 μm. Enlargements represent a single z slice; scale bars: 2 μm.

This hypothesis was tested by treating infected cells with the small molecule PF74, which acts as a competitive inhibitor of CA-CPSF6 interaction (Bejarano et al., 2019) and may thus displace CPSF6 from the subviral complex. CD4^+^ T cells infected with NNHIV ANCH for 5 h or 22 h were treated with PF74 or solvent for 2 h before fixation and IF staining (Figure 6b-d). The CPSF6 signal was completely lost from nuclear HIV-1 subviral complexes upon PF74 treatment at both time points, while IN-FP punctae stayed intact (Figure 6b, lower panel, c). PF74-mediated removal of CPSF6 from the nuclear subviral complex also revealed a strong CA IF signal, which was absent in solvent treated cells (Figure 6e). No difference was observed for CA IF signals on extracellular particles and on cytoplasmic or nuclear envelope-associated subviral complexes upon PF74 treatment, while the nuclear CA signal became strongly enhanced (Figure 6f, g). Thus, the failure to detect nuclear CA by IF in CD4^+^ T cells is due to shielding of epitopes by the accumulation of CPSF6 and not due to CA being lost upon nuclear entry in these cells. Compared to extracellular virions, subviral particles in the cytosol and at the nuclear envelope displayed a reduced CA signal due to loss of free CA molecules from the post-fusion complex. This phenotype was most notable for nuclear envelope associated complexes at later time points whose CA signal corresponded to ~ 50 % of that of complete particles (Figure 6g). Of note, nuclear subviral complexes showed a higher mean average CA signal compared to complete virions, supporting nuclear clustering of CA containing complexes; this was most evident at earlier time points (Figure 6f, g). After PF74 treatment we could also observe some nuclear eGFP.OR3 punctae representing HIV-1 cDNA that were associated with IN-FP and clearly CA positive, with a strong signal (Figure 6h). Upon separation from eGFP.OR3 punctae, CPSF6 clusters remained associated with the IN-FP, confirming that the viral cDNA separates from an IN-FP/CA/CPSF6 positive nuclear structure in the nucleus of infected primary CD4^+^ T cells as well (Figure 6i).

Finally, we adapted the ANCHOR system to primary MDM by transducing these cells with an eGFP.OR3 expressing lentiviral vector. Three days post transduction, MDM were infected with VSV-G pseudotyped NNHIV ANCH containing IN.mScarlet for 24 h. A similar phenotype as shown above for TZM-bl cells and primary CD4^+^ T cells was observed in MDM: the majority of eGFP.O3 punctae was detected separated from, but often in close vicinity of IN.mScarlet punctae (Figure 6j).

## Discussion

In this study, we adapted a single molecule labeling method to study the dynamics of HIV-1 cDNA in living cells. Using this system, we showed that the HIV-1 ANCH dsDNA recognizing OR3 marker is only recruited to the viral cDNA inside the nucleus, while no OR3 punctae were observed associated with viral structures in the cytosol despite the presence of abundant reverse transcription products. Both, integrated and unintegrated HIV-1 cDNA were detected by OR3 in the nucleus. Metabolic labeling of nascent DNA revealed that nuclear eGFP.OR3 punctae contained significantly higher DNA amounts compared to cytoplasmic or nuclear subviral complexes lacking the eGFP.OR3 signal. Together with recent reports employing indirect RT inhibitor time-of-addition assays, which showed that reverse transcription remains sensitive to inhibition until after nuclear import (Burdick et al., 2020; Dharan et al., 2020), and an elegant experiment showing that positive and negative strand-specific HIV DNA hybridisation probes only co-localize inside the nucleus (Dharan et al., 2020), our data support completion of HIV-1 reverse transcription in the nucleus. Furthermore, HIV-1 cDNA separated from an IN fusion protein (IN-FP), often used as a marker for the HIV-1 replication complex, inside the nucleus; IN fusions are thus not suitable for tracking HIV-1 cDNA in the nucleus. We expect that unfused IN, also present in the replication complex, will remain - at least partially - with the cDNA to mediate chromosomal integration and thus separates from the bulk of IN-FP. Nuclear IN-FP punctae were strongly positive for the viral CA protein and CA, CPSF6 and IN-FP signals stayed together after separation from the viral cDNA.

OR3 recruitment to the HIV-1 cDNA requires the dsDNA to be accessible to the 66 kDa eGFP.OR3 fusion protein, which depends on loss of integrity of the capsid shell. CA signal intensity on nuclear IN-FP positive and eGFP.OR3-negative structures was equal to or higher than observed for cytoplasmic complexes, suggesting that the bulk of CA stays associated with the viral replication complex and the viral DNA remains encased inside a closed capsid or capsid-like structure until after nuclear entry. Using CLEM-tomography of IN-positive and eGFP.OR3-negative nuclear complexes, we observed morphologically intact cone-shaped structures with internal density representing the nucleoprotein complex, which closely resembled HIV-1 capsids inside the cytosol or in complete virions. This finding is consistent with our recent study showing that the nuclear pore channel is sufficiently large to accommodate the HIV-1 core and apparently intact cone-shaped HIV-1 capsids can enter the nucleus through intact nuclear pores (Zila et al., 2020). Taken together, these results indicate that reverse transcription initiates in the cytoplasm inside a complete or largely complete capsid, and this capsid-encased complex trafficks into the nucleus, where reverse transcription is completed; subsequently it must be uncoated for integration to occur. Formation of eGFP.OR3 punctae requires both, completion of reverse transcription and - at least partial - uncoating, and these two events may conceivably occur in a coordinated manner.

IN-FP (and CA) positive nuclear complexes at later time points were often observed in close vicinity but clearly separated from eGFP.OR3 punctae, and live cell microscopy confirmed separation of the two markers from a single focus over time. Both markers retained their focal appearance and could thus be analysed by CLEM. Electron tomography of late IN-FP positive complexes revealed electron-dense structures that resembled broken HIV-1 capsids or capsid-like remnants lacking the density of the nucleoprotein complex. In contrast, eGFP.OR3-positive and IN-negative nuclear subviral structures never exhibited an electron-dense lining resembling the capsid shell, and these complexes were always negative for CA by immunofluorescence. These results indicate that viral cDNA associated with some replication proteins emanates from the broken capsid, which retains the bulk of CA and capsid-associated proteins. Uncoating therefore does not appear to occur by cooperative disassembly of the CA lattice, but by physically breaking the capsid shell and loss of irregular capsid segments. We often observed clustering of capsids or capsid-remnants inside the nucleus indicating preferred trafficking routes of subviral complexes.

The broken capsid-remnant structures inside the nucleus of infected cells closely resembled ruptured HIV-1 cores observed in a recent study analyzing HIV-1 cDNA formation and integration in an *in vitro* system using purified viral cores (Christensen et al., 2020). These authors reported partially broken capsid shells with irregular defects at time points when endogenous cDNA formation was largely completed; they also detected polynucleotide loops emanating from the holes in the capsid lattice. Theoretical models and AFM-studies had suggested that the volume of double-stranded HIV-1 b-DNA cannot be accommodated inside the intact capsid and that the resulting pressure might mechanically trigger uncoating (Rankovic et al., 2017; Rouzina & Bruinsma, 2014). Taken together, these results suggest that the growing dsDNA inside the viral capsid in the nucleus may eventually lead to local rupture of the capsid lattice, concomitantly allowing completion of reverse transcription and triggering uncoating of the proviral DNA. It must be kept in mind, however, that lentiviral vectors with much shorter length of the vector RNA efficiently transduce cells, and this will need to be analyzed in future studies. Nuclear import is not required for completion of cDNA synthesis and loss of capsid integrity since similar structures were detected in the *in vitro* system (Christensen et al., 2020). The observation that the viral cDNA was not fully released from the viral core *in vitro* suggests, however, that the nuclear environment may play a role in this process. The described pathway appears to be conserved in HeLa reporter cells and primary HIV-1 sensitive CD4^+^ T-cells and MDM: separation of IN/CA complexes from the OR3-positive cDNA inside the nucleus of infected cells was observed in all cell types, and the IN-positive subviral complexes exhibited a strong CA signal in all cases. Previous failure to detect CA on nuclear complexes in T cells has been due to epitope masking by the cellular CPSF6 protein and the current results thus indicate a common pathway for early HIV-1 replication in different cell types including primary target cells of HIV-1 infection.

STED and CLEM analysis revealed elongated structures with regularly spaced globular densities at the position of eGFP.OR3-positive punctae that had separated from the IN fusion protein and CA. These structures resembled chromatinized DNA (Miron et al., 2020), in line with biochemical evidence that HIV-1 cDNA is rapidly chromatinized when it becomes accessible to the nucleoplasm (Geis & Goff, 2019). Detection of a cone-shaped structure lacking the electron-dense internal nucleoprotein signal and directly associated with an elongated chromatin-like structure at a position that was positive for both IN-FP and eGFP.OR3 may have captured a subviral complex in the process of uncoating. We suggest that chromatinization of HIV-1 cDNA emerging from the broken capsid may facilitate complete uncoating of the genome, which could explain why viral cDNA remained largely associated with the capsid structure in the *in vitro* system.

In conclusion, our results indicate that complete or largely complete HIV-1 capsids enter the nucleus of infected cells, where reverse transcription is completed and the viral cDNA genome is released by physical disruption rather than by cooperative disassembly of the capsid lattice (Figure 7). The viral capsid thus plays an active role in the entire early phase of HIV-1 replication up to chromosomal integration and appears to be important for cytoplasmic trafficking, reverse transcription, shielding of viral nucleic acid from the innate immune system, nuclear entry and intranuclear trafficking. The cone-shaped HIV-1 capsid with its fullerene geometry thus is the key orchestrator of early HIV-1 replication.

**Figure 7.**
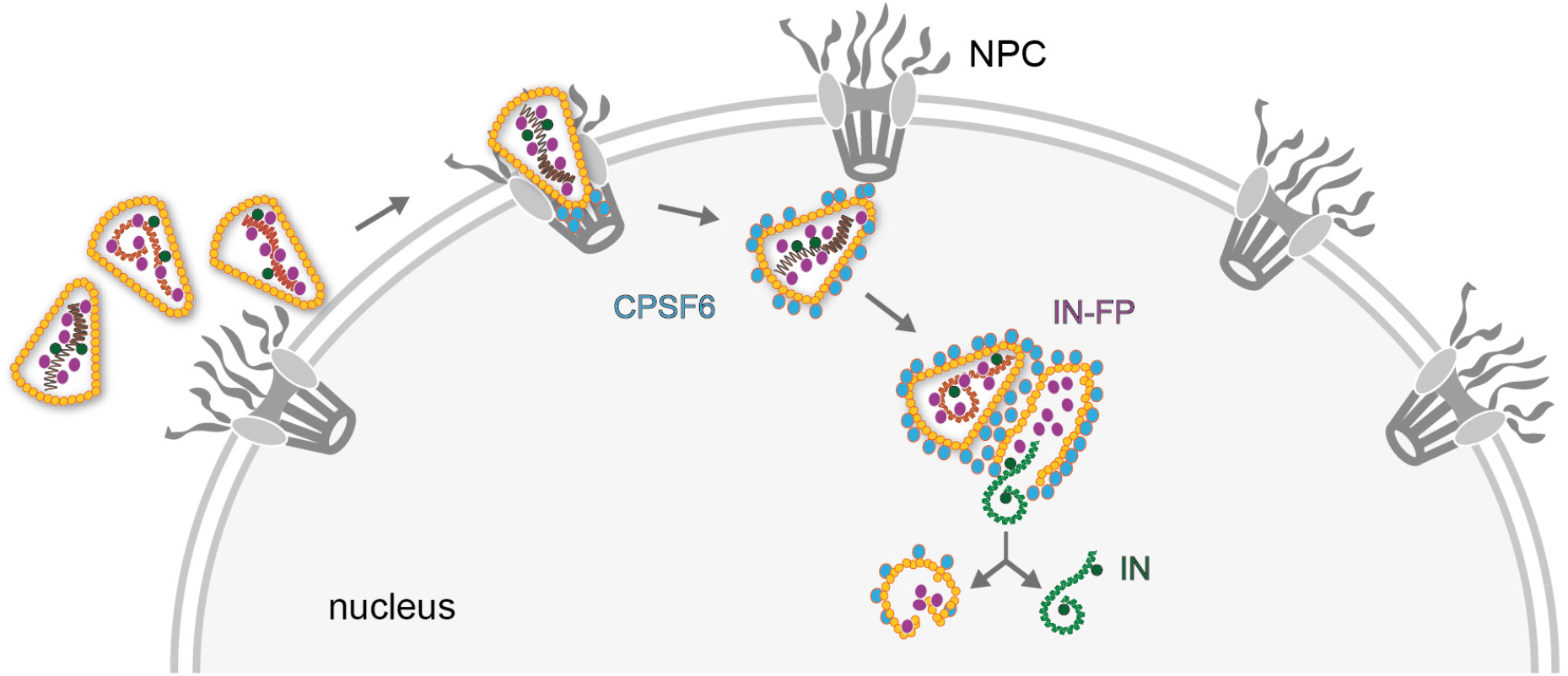
Model of HIV-1 nuclear entry and uncoating. Apparently intact HIV capsids are imported into the nucleus through nuclear pore complexes (NPC) retaining their cone-shaped morphology. CPSF6 releases the cores from the NPC and clusters on nuclear capsids. Multiple capsids accumulate at certain positions within the nucleus, and plus strand synthesis of the viral double-stranded cDNA is completed in the nucleus. Physical disruption of the capsid releases the completed cDNA into the nucleoplasm, where it becomes integrated into the host cell genome in the vicinity of the uncoating site. Empty remnants of the broken capsid, associated with incorporated viral proteins that are not part of the cDNA complex, remain as distinct structures in the nucleus for prolonged times after uncoating.

## Materials and Methods

### Key resources table

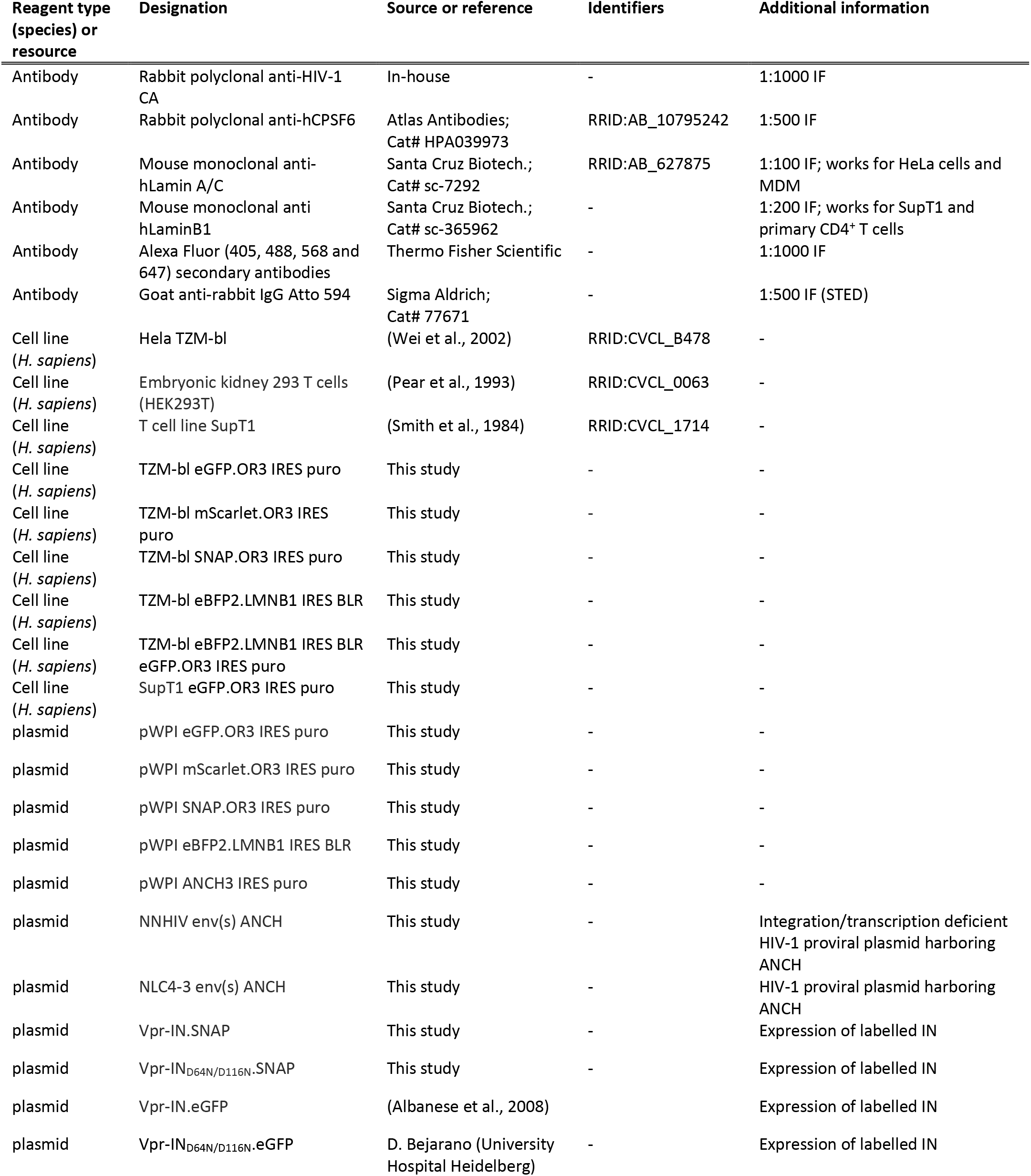
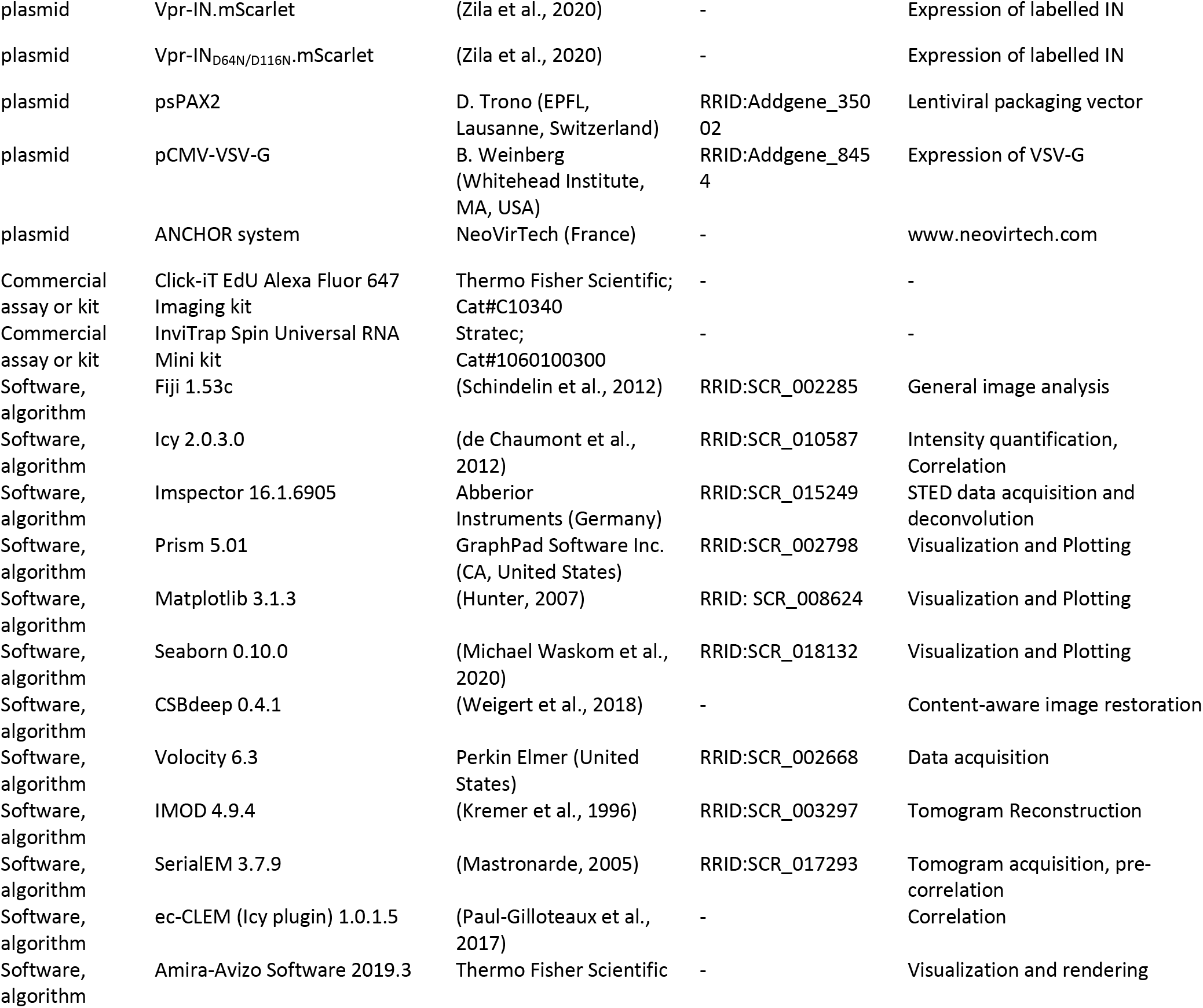

### List of primers

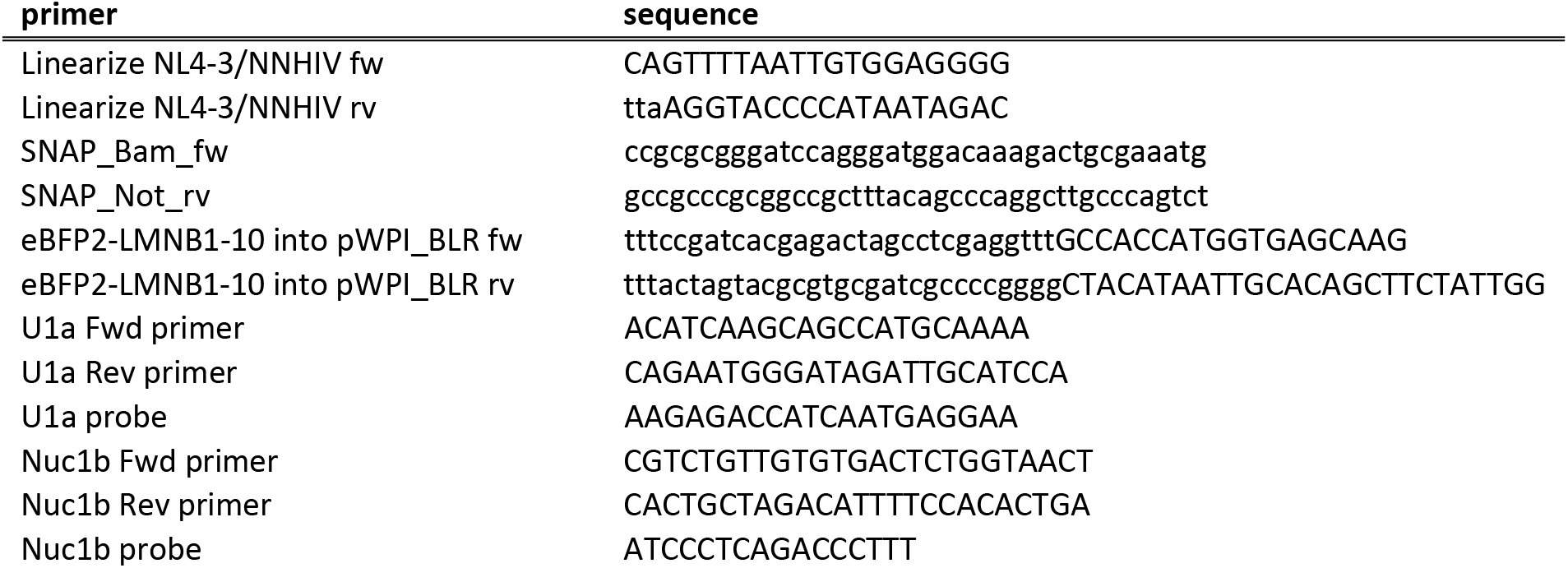
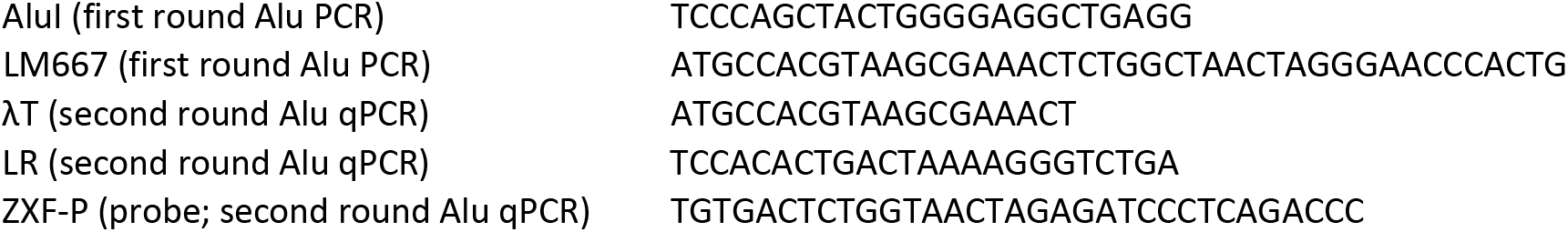

### Plasmids

Plasmids were cloned using standard molecular biology techniques and verified by commercial Sanger sequencing (Eurofins Genomics, Germany). Gibson assembly was performed using the NEB HiFi Mastermix (New England Biolabs, USA) and 30 bp overlap regions. PCR was performed using Q5 High-Fidelity DNA Polymerase (New England Biolabs) according to manufacturer’s instructions with primers purchased from Eurofins Genomics. E. coli DH5α and Stbl2 (Trinh et al., 1994, p. 2) (Thermo Fisher Scientific, USA) were used for amplification of standard plasmids or LTR containing plasmids, respectively.

#### Derivatives of pNL4-3 and pNNHIV harboring ANCH1000 within the env gene

To facilitate the cloning procedure, EcoRI/XhoI fragments comprising the env region of HIV were subcloned from pNLC4-3 (Bohne & Kräusslich, 2004) and its non-replication competent derivative pNNHIV (Zila et al., 2020) into pcDNA3.1^(+)^ (Thermo Fisher Scientific). These constructs were PCR linearized, deleting a ~ 1000 bp region (nt 130-1113) within the *env* coding sequence. The ANCH3 1000 bp sequence was PCR amplified from pANCH3 (NeoVirTech, France), introducing a stop codon directly upstream of ANCH3, and transferred into the linearized vector fragments using Gibson assembly. The modified fragments were transferred into pNL4-3 or pNNHIV backbones using EcoRI/XhoI.

#### Vpr-IN.SNAP and Vpr-IN_D64N/D116N_.SNAP

The SNAP-tag gene was PCR amplified from pSNAP-tag(m) (Addgene #101135) and cloned into pVpr-IN.eGFP(Albanese et al., 2008) using BamHI/NotI, substituting the eGFP gene for the SNAP-tag coding region. To generate the Vpr-IN_D64N/D116N_.SNAP mutant, the IN_D64N/D116N_ from Vpr-IN_D64N/D116N_.eGFP (gift from D. A. Bejarano) was cloned into Vpr-IN.SNAP using BamHI/NotI.

#### Lentiviral transfer vectors harboring the ANCH sequence and eGFP.OR3, SNAP.OR3, mScarlet.OR3 and eBFP2.LMNB1 coding sequences

The ANCH3 1000 bp sequence was PCR amplified from pANCH3 (NeoVirTech) and cloned by Gibson assembly into pWPI IRES puro (Trotard et al., 2016) linearized with NotI. The eGFP.OR3 gene was PCR amplified from peGFP-OR3 (NeoVirTech) and transferred via Gibson assembly into PmeI/BamHI linearized pWPI IRES puro to create the expression cassette EF1-alpha eGFP.OR3 IRES puro. The SNAP gene was amplified from pVpr.IN.SNAP and the mScarlet (WT) (Bindels et al., 2016) gene from the mScarlet C1 vector (Addgene #85042) and placed N-terminal to OR3 into PCR linearized pWPI EF1-alpha OR3 IRES puro backbone by Gibson assembly, substituting eGFP. The eBFP2.LMNB1 gene was amplified from pEBFP2-LaminB1-10 (Addgene #55244) and transferred via Gibson assembly into PmeI/BamHI linearized pWPI IRES BLR (Trotard et al., 2016).

### Cell culture

HEK293T (Pear et al., 1993) (RRID: <u>CVCL_0063</u>), HeLa TZM-bl (Wei et al., 2002) (RRID: <u>CVCL_B478</u>) and SupT1 (Smith et al., 1984) (RRID: <u>CVCL_1714</u>) cell lines were authenticated using STR profiling (Eurofins Genomics) and monitored for mycoplasma contamination using the MycoAlert mycoplasma detection kit (Lonza Rockland, USA). Cells were cultured at 37 °C and 5 % CO_2_ in Dulbecco’s Modified Eagle’s Medium (DMEM; Thermo Fisher Scientific) containing 4.5 g l^−1^ D-glucose and L-glutamine supplemented with 10 % fetal calf serum (FCS; Sigma Aldrich, USA), 100 U ml^−1^ penicillin and 100 μg ml^−1^ streptomycin (PAN Biotech, Germany) (adherent cell lines) or in RPMI 1640 (Thermo Fisher Scientific) containing L-glutamine supplemented with 10 % FCS, 100 U ml^−1^ penicillin and 100 μg ml^−1^ streptomycin (SupT1 cells). Primary CD4^+^ T cells were cultured in RPMI 1640 containing L-glutamine supplemented with 10 % heat-inactivated FCS, 100 U ml^−1^ penicillin and 100 μg ml^−1^ streptomycin. Monocyte-derived macrophages (MDM) were cultured in RPMI 1640 containing 10 % heat-inactivated FCS, 100 U ml^−1^ penicillin, 100 μg ml^−1^ streptomycin and 5 % human AB serum (Sigma Aldrich).

*Isolation of primary cells.* CD4^+^ T cells were enriched from blood of healthy donors using RosetteSep Human CD4^+^ T cell enrichment cocktail (Stemcell Technologies, Canada) according to the manufacturer’s instructions followed by Ficoll density gradient centrifugation. Subsequently, cells were activated using human T-Activator CD3/CD28 Dynabeads (Thermo Fisher Scientific) and 90 U/ml IL-2 for 48-72 h. MDMs were isolated from buffy coats of healthy blood donors as described previously (Bejarano et al., 2019).

### Generation of cell lines

Lentiviral vector particles were produced by co-transfection of packaging plasmid psPAX2 (Addgene #12260), the respective lentiviral transfer vector pWPI, the envelope expression plasmid pCMV-VSV-G (Addgene #8454) and pAdvantage (Promega, USA) in a ratio of 1.5: 1.0: 0.5: 0.2 μg into HEK293T cells (4×10^5^ cells/well seeded the day before in 6 well plates) using polyethylenimin (PEI; 3 μl of 1 mg/ml PEI per μg DNA). At 48 h post transfection the tissue culture supernatant was harvested and filtered through 0.45 μm mixed cellulose ester (MCE) filters. SupT1 (1 ml of freshly 1:4 diluted cells) or TZM-bl (5×10^4^ cells/well seeded the day before in 12 well plates) cells were transduced using 50-500 μl supernatant. At 48 h post transduction selection with 1 μg/ml puromycin or 5 μg/ml blasticidin was initiated. For transduction of MDM, lentiviral vectors were produced with Vpx_mac239_ (Bejarano et al., 2018) by calcium phosphate transfection of packaging plasmid pΔR8.9 NSDP(Pertel et al., 2011), containing a Vpx interaction motif in Gag, pWPI eGFP.OR3 IRES puro, Vpx expression plasmid pcDNA.Vpxmac239 (Sunseri et al., 2011) and pCMV-VSV-G at a ratio of 1.33: 1.00: 0.17: 0.33 μg (68 μg / T175 flask). MDM were differentiated in human AB serum (Sigma Aldrich) from monocytes (Bejarano et al., 2019) in 15-well μ-Slides Angiogenesis (ibidi, Germany) for 10 d and transduction was performed 2 d prior to infection.

### Production of viral particle stocks

pNLC4-3 or pNNHIV ANCH, a Vpr-IN plasmid (Vpr-(SNAP/eGFP/mScarlet).IN or Vpr-(SNAP/eGFP/mScarlet).IN_D64N/D116N_) and pCMV-VSV-G or pCAGGS.NL4-3-Xba (Bozek et al., 2012) were transfected in a ratio of 7.7: 1.3: 1.0 μg into HEK293T cells using calcium phosphate (70 μg / T175 flask). Medium was changed at 6-8 h and cells were further incubated for 48 h. Supernatant was harvested and filtered through 0.45 μm MCE before ultracentrifugation through a 20 % (w/w) sucrose cushion (2 h, 107.000 g). Pellets were resuspended in phosphate-buffered saline (PBS) containing 10 % FCS and 10 mM HEPES (pH 7.5), and stored in aliquots at −80 °C. Virus was quantified using the SYBR Green based Product Enhanced Reverse Transcription assay (SG-PERT) (Pizzato et al., 2009). MOI of infectious particles was determined by titration on TZM-bl cells and immunofluorescence staining against HIV CA at 48 h p.i‥ The proportion of positive cells was counted in > 10 randomly selected fields of view.

### Labeling of SNAP-tagged virus and infection

3.33 × 10^3^ TZM-bl cells were seeded into 15-well μ-Slides Angiogenesis (ibidi) the day before infection. Stock solutions of SNAP-Cell® TMR-Star or SNAP-Cell® 647-SiR (New England Biolabs) in DMSO were diluted to 4 μM in complete medium, mixed 1:1 with IN.SNAP particles and incubated at 37 °C for 30 min. Labelled particles were added to cells at 5-30 μUnits RT /cell in 50 μl. For VSV-G pseudotyped pNL4-3 ANCH, 30 μUnits RT per TZM-bl cell corresponds to ~ MOI 6 in TZM-bl cells. Infection of MDM was performed with NNHIV ANCH (50 μl, 120 μUnits RT/cell). Infection of suspension cells was performed with 2×10^4^ cells per 15 well μ-Slide in 96 well v-bottom plates (40 μl; 30 μU RT/cell). For PF-3450074 (PF74; Sigma Aldrich) experiments in primary CD4^+^ T cells, medium was changed at 5 or 22 h to medium containing 15 μM PF74 or DMSO, for 1 h before transfer to PEI coated (with 1 mg/ml PEI for 60 min) μ-Slides. Slides were incubated for 1 h for cell attachment prior to fixation. Efavirenz (EFV; Sigma Aldrich), Raltegravir (Ral; AIDS Research and Reference Reagent Program, Division AIDS, NIAID) and Azidothymidine (AZT) were added at time of infection. Flavopiridol (Sigma Aldrich) and 5,6-dichloro-1-beta-D-ribofuranosylbenzimidazole (DRB; Sigma Aldrich) were added 8 h prior to fixation or RNA extraction. 10 μM EdU (Thermo Fisher Scientific) and 6 μM APC (Sigma Aldrich) were added at the time of infection.

### Fixation immunofluorescence staining and EdU click-labeling

Samples were washed with PBS and fixed (15 min, 4 % PFA), washed again three times using PBS, permeabilized with 0.5 % Triton X-100 (TX-100) for 10 or 20 min and washed again. In indicated experiments, cells were extracted using ice-cold methanol for 10 min. Afterwards, cells were washed two times using 3 % bovine serum albumin (BSA)/PBS and blocked for 30 min with 3 % BSA. Primary antibody in 0.5 % BSA was added for 1 h at room temperature. After washing three times with 3 % BSA/PBS, secondary antibody in 0.5 % BSA was added for 1 h at room temperature and samples were washed and stored in PBS. For EdU incorporation experiments, cells were click-labeled for 30 min at room temperature using the Click-iT EdU-Alexa Fluor 647 Imaging Kit (Thermo Fisher Scientific) according to the manufacturer’s instructions.

### DNA fluorescent in situ hybridization (FISH)

Biotinylated HIV-1 FISH probes were prepared with the Nick Translation Kit (Roche, Germany) according to the manufacturer’s instructions. Probes were purified with Illustra Microspin G-25 columns (GE Healthcare, UK) according to the manufacturer’s instructions and ethanol precipitated with human Cot-1 DNA (Roche, Germany) and herring sperm DNA (Sigma Aldrich). Probes were resuspended in 10 μL formamide, incubated at 37 °C for 15-20 min and 10 μL of 20 % dextran/4X saline-sodium citrate (SSC) buffer was added.

1.25 × 10^4^ TZM-bl cells/well were seeded on PEI (1 mg/ml) coated glass cover slips and infected with VSV-G pseudotyped IN.SNAP.SiR labelled NNHIV ANCH (30 μU/cell). At 24 h cells were fixed for 10 min with 4% PFA/PBS, permeabilized with 0.5 % TX-100/0.1 % Tween/PBS for 10 min at room temperature, and washed in 0.1 % Tween/PBS. Following 30 min blocking in 4 % BSA/PBS, cells were incubated with rabbit anti-GFP antibody (ab6556; Abcam, UK), diluted (1:2000) in 1 % BSA/PBS overnight at 4 °C. Cells were washed in 0.1 % Tween/PBS and incubated with secondary Alexa Fluor antibody (Thermo Fisher Scientific) for 1 h at room temperature. Cells were fixed for 10 min with 0.5 mM ethylene glycol bis(succinimidyl succinate) (EGS)/PBS, washed in 0.1 % Tween/PBS and permeabilized with 0.5 % TX-100/0.5 % saponin/PBS for 10 min. Cells were incubated for 45 min in 20 % glycerol/PBS and subjected to four glycerol/liquid N_2_ freeze-thaw cycles. Samples were rinsed, incubated in 0.1 M HCl for 10 min, equilibrated in 2X SSC for 20 min and left in hybridization buffer (50 % Formamide/2X SSC) for 30 min. Samples were then washed in PBS, treated with 0.01 N HCl/0.002 % pepsin (3 min, 37 °C) and quenched by addition of 1X PBS/1 M MgCl_2_. Fixation with 4 % PFA/PBS and PBS wash was followed by treatment with 100 μg/ml RNase A (PureLink, Invitrogen, USA) in 2x SSC for 30 min at 37 °C, washing and overnight storage in hybridization buffer.

1-10 μL of heat-denatured FISH probe (7 minutes at 95 °C) was loaded onto glass slides covered with coverslips coated with prepared cells. Slides were sealed in a metal chamber heated at 80 °C for 7 min, and incubated for 48 h at 37 °C. Samples were washed in 2X SSC at 37 °C, followed by 3 washes in 0.5 X SSC at 56 °C. FISH detection was performed using anti-biotin antibody (SA-HRP) and a FITC/Cy5 coupled secondary antibody with the TSA Plus system (Perkin Elmer, USA). Coverslips were stained with Hoechst, mounted on glass slides and imaged using the Nikon/Andor SDCM system described below.

### Confocal microscopy

Spinning disc confocal microscopy (SDCM) was performed on an inverted Nikon Eclipse Ti2 (Nikon, Japan) microscope equipped with a Yokogawa CSU-W1 Spinning Disk Unit (Andor, Oxford Instruments, United Kingdom) and an incubation chamber (37 °C, 5 % CO_2_). Imaging was performed using a 100 × oil-immersion objective (Nikon CFI Apochromat TIRF 100X Oil NA 1.49) and either single or dual channel EMCCD camera setup (ANDOR iXon DU-888) recording the eBFP2 (405/420-460), eGFP (488/510-540 nm), mScarlet (568/589-625 nm) and SiR (647/665-705 nm) channels with a pixel size of 0.13 μm. 3D stacks of 10-30 randomly chosen positions were automatically recorded with a z-spacing of 0.3-0.5 μm using the Nikon Imaging Software Elements v5.02. For CLEM experiments a Perkin Elmer Ultra VIEW VoX 3D spinning disk confocal microscope (Perkin Elmer, United States) with a 100 x oil immersion objective (NA 1.4; Perkin Elmer) was used.

### Live cell imaging

Medium was exchanged for 50 μl imaging medium containing FluoroBrite DMEM (Thermo Fisher Scientific), 10 % FCS, 4 mM GlutaMAX (Gibco Life Technologies), 2 mM sodium pyruvate (Gibco Life Technologies), 20 mM HEPES pH 7.4, 100 U/ml Penicillin and 100 μg/ml Streptomycin (PAN-Biotech). Samples were transferred to the SDCM setup described above. 3D stacks were recorded up to 24 h (time interval of 3-30 min, z-spacing 0.5 μm). Data from Figure 1g and f was fit to a four-parametric logistic population growth model with variable slope using Prism 5.01 (Graphpad)

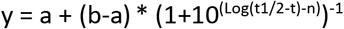

with y = normalized eGFP.OR3 events per cell, a = Y value at the bottom plateau, b = Y value at the top plateau, t = h p.i., t_1/2_ = time at half maximal signal and n = slope factor.

### STED microscopy

3.33 × 10^3^ SNAP.OR3 expressing cells were seeded on 15-well μ-Slides Angiogenesis (ibidi) and infected as described above. Prior to fixation, cells were incubated with 2 μM SNAP-Cell® 647-SiR (New England Biolabs) for 30 min at 37 °C, washed three times and fixed with 4 % PFA (15 min). STED microscopy was performed using a 775 nm STED system (Abberior Instruments GmbH, Germany) equipped with a 100 x oil immersion objective (NA 1.4; Olympus UPlanSApo). STED Images were acquired using the 590 and 640 nm excitation laser lines while the 405 and 488 laser lines were acquired in confocal mode. Nominal STED laser power was set to 80 % of the maximal power (1250 mW) with 20 μs pixel dwell time and 15 nm pixel size. STED Images were linearly deconvolved with a Lorentzian function (FWHM 50 nm) using the Richardson-Lucy algorithm and the software Imspector (Abberior Instruments GmbH).

### Image analysis and data visualization

The images were filtered in Fiji/ImageJ (Schindelin et al., 2012) with a mean filter (kernel size: 0.26 × 0.26 μm) to reduce noise. For visualization some of the low signal-to-noise 3D movies were denoised using content-aware image restoration (CARE) (Weigert et al., 2018) as indicated. Convolutional neuronal networks were trained with a set of fixed cell images recorded with high laser power/long camera exposure (ground truth) and low laser power/short camera exposure (noisy training input) using the Python toolbox CSBdeep (Weigert et al., 2018). The model was then applied to reconstruct raw movies in Fiji using the CSBdeep CARE plugin. Quantification of fluorescent spot intensities was performed using Icy (de Chaumont et al., 2012). Raw 3D stacks were used to detect volumes of IN objects using the spot detector plugin. Methanol-induced shrinkage of cells in z orientation and infection-induced invaginations of the nuclear envelope renders automated nuclei detection difficult. To ensure that only truly nuclear objects (and not still NE-associated ones) were classified as such, they were manually curated and ambiguous particles were excluded. Objects displaying positive signals in the lamin channel were classified as nuclear envelope (NE) associated in the TZM-bl experiments. For measurements in primary CD4^+^ T cells we applied a more stringent classification, manually excluding objects that did not colocalize with lamin in the major part of the signal or were localized above/below the focal planes of the nucleus. Cell-specific local background was subtracted for CA quantification. For quantification of CPSF6 intensities, the diffusive nuclear expression level of the cell was measured (using a ROI without punctae) and intensities of CPSF6 accumulations at IN objects were normalized to the expression level of the cell. Nuclear OR3 punctae were counted if their intensity was ≥ 20 % above the diffuse nuclear expression level of the respective cell. Colocalization was scored when the pixel areas of the respective fluorescent spots (partially) overlapped. Tracking was performed using the Fiji plugin Manual Tracking. Fiji standard “Fire” lookup table (LUT) was used for visualization of single channel images. Statistical tests were performed using Prism v5.01 (GraphPad Software Inc., USA). Data were plotted using Prism v5.01 or the Python statistical data visualization package matplotlib v3.1.3 (Hunter, 2007) and seaborn v0.10.0 (Michael Waskom et al., 2020). Graphs show mean with error bars defined in the figure legends.

### Quantification of RT products

Particle preparations filtered through 0.45 μm CME were treated with 15 U/mL DNAseI (Sigma Aldrich) / 10 mM MgCl_2_ for 3-4 h at 37 °C prior to ultracentrifugation. 5×10^4^ TZM-bl cells were seeded into 24-well plates and infected the following day using 10-30 μU RT/cell. At the indicated h p.i. cells were washed, scraped and lysed using 50 μl of 10 mM Tris-HCl pH 9.0, 0.1 % TX-100, 400 μg/mL proteinase K (Thermo Fisher Scientific) at 55 °C overnight. Proteinase K was inactivated at 95 °C for 10 min and lysates were stored at −20 °C. Alternatively, DNA was purified from cells using the DNeasy Blood and Tissue Kit (Qiagen, Germany) according to the manufacturer’s instructions. Lysates were directly used as input for ddPCR; purified DNA was prediluted to ~20 ng/μl. For *gag* cDNA detection, this input was additionally diluted 1:20 to prevent saturation. ddPCR was performed using the QX200 droplet generator/reader (BioRad, USA) and analyzed using QuantaSoft v1.7.4 (BioRad) as described earlier (Bejarano et al., 2018; Zila et al., 2019).

### Quantification of viral RNA transcripts

eBFP2.LMNB1 and eGFP.OR3 expressing TZM-bl cells were infected with VSV-G pseudotyped NL4-3 ANCH (5 μUnits RT/cell; MOI ~1) for 55 h. 20 μm RAL was added at the time of infection. 8 h prior to RNA extraction and purification using the Invitrap spin universal RNA mini kit (Stratec biomedical, Germany), medium was changed for medium containing 1-25 μM flavopiridol (Sigma Aldrich) or 1-25 μM 5,6-dichloro-1-beta-D-ribofuranosylbenzimidazole (DRB; Sigma Aldrich) as indicated. Quantitative reverse transcription PCR was performed as previously described (Marini et al., 2015). Briefly, messenger RNA (mRNA) levels were quantified by TaqMan quantitative RT-PCR (qRT-PCR). First, the RT reaction was performed using M-MLV RT (Invitrogen) and a random primer set (Invitrogen, cat. no.: 48190011), followed by qPCR using HIV-1 primers and probes (specific for transcription of the first nucleosome nuc1a or gag/u1a) and the housekeeping genes 18S and GAPDH (both containing VIC&trade/TAMRA&trade fluorescent probe; Applied Biosystems, USA) as controls (see list of primers).

### Quantification of integrated proviral DNA using Alu PCR

2×10^5^ SupT1 cells were infected using VSV-G pseudotyped HIV-1 NL4-3 ANCH (10 μU RT/cell) for 24 or 48h. 20 μm Ral or 20 μm EFV was added at the time of infection. Cells were washed, lysed and genomic DNA was extracted using the DNeasy Blood and Tissue Kit (Qiagen) according to the manufacturer’s instructions. Nested Alu-LTR PCR was performed as described before (Tan et al., 2006). Briefly, Alu-LTR fragments were amplified starting from 100 ng of genomic DNA. The product of the first amplification was diluted 1:50 in H_2_O and amplified by qPCR. The B13 region of the housekeeping gene lamin B2 was amplified from 10 ng of genomic DNA for normalization (Livak & Schmittgen, 2001). The copy number (Log10) of integrated HIV-1 DNA per million cells was calculated using a standard curve obtained by serially diluting DNA from HIV-1 infected and sorted p24^+^ cells with DNA from uninfected cells as described before (Shytaj et al., 2020).

### CLEM sample preparation

1.2×10^5^ TZM-bl mScarlet.OR3 cells were grown on 3-mm sapphire discs in a 35 mm glass-bottom dish (MatTek, USA). Cells were infected with VSV-G pseudotyped IN.SNAP.SiR labelled NNHIV ANCH (30 μU RT/cell) and incubated for 24 h at 37 °C. Subsequently, cells were cryo-immobilized by high pressure freezing (HPF) (HPM010; Abra Fluid, Switzerland) and transferred to freeze-substitution (FS) medium (0.1% uranyl acetate, 2.3 % methanol and 1 % H_2_O in Acetone) tempered at −90 °C. Freeze-substituted samples were embedded in Lowicryl HM20 resin (Polysciences, USA) inside a FS device (AFS2, Leica, Germany) equipped with a robotic solution handler (FSP, Leica). FS and embedding into Lowicryl resin was performed according to Kukulski *et al.* (Kukulski et al., 2011) with modifications (Zila et al., 2020). Temperature was raised to −45 °C at 7.5 °C/h. Samples were washed with acetone (4 × 25 min) and infiltrated with increasing concentrations of Lowicryl in acetone (25, 50 and 75 %; 3 h each), while raising temperature to −25 °C (3.3 °C / h). The acetone-resin mixture was replaced by Lowicryl HM20 for 1 h and the resin was exchanged three times (3, 5 and 12 h). Samples were polymerized under UV light for 24 h at −25 °C. Polymerization continued for an additional 24 h while the temperature was raised to 20 °C at 3.7 °C/h.

### CLEM and electron tomography (ET)

Thick resin sections (250 nm) were cut using a microtome (EM UC7, Leica) and placed on a slot (1 × 2 mm) EM copper grid covered with a formvar film (FF2010-Cu, Electron Microscopy Sciences, USA). Sections were covered by 0.1 μm TetraSpeck microsphere fiducials (Thermo Fisher Scientific). Nuclear regions were stained with 50 μg/ml Hoechst (Thermo Fisher Scientific). For SDCM, grids were transferred to 25 mm glass coverslips mounted in a water-filled ring holder (Attofluor cell chamber, Thermo Fisher Scientific). Z stacks of cell sections were acquired using the PerkinElmer UltraVIEW VoX 3D Spinning-disc Confocal Microscope described above (*z-*spacing 200 nm). To identify mScarlet.OR3 and IN.SNAP.SiR signals in cell sections, images were visually examined using Fiji (Schindelin et al., 2012). Subsequently, both sides of EM grids were decorated with 15 nm protein-A gold particles for tomogram alignment and contrasted with 3 % uranyl acetate (in 70 % methanol) and lead citrate. Individual grids were placed in a high-tilt holder (Fischione Model 2040) and loaded into the Tecnai TF20 (FEI, Eindhoven, Netherlands) electron microscope operated at 200 kV, equipped with a field emission gun and a 4 K by 4 K pixel Eagle CCD camera (FEI, USA). To identify positions for ET, a full grid map was acquired using SerialEM (Mastronarde, 2005) and acquired electron micrographs were pre-correlated with imported SDCM images in SerialEM using fiducials as landmark points (Schorb et al., 2017). Single-axis electron tomograms of selected regions were then carried out. Tomographic tilt ranges were from −60 ° to 60 ° with an angular increment of 1 ° and pixel size 1.13 nm. Alignments and 3D reconstructions of tomograms were done using IMOD (Kremer et al., 1996). For high precision post-correlation, tomograms of cell sections were acquired at lower magnification with 4 ° increment and 6.3 nm pixel size. Post-correlation was performed using the eC-CLEM plugin (Paul-Gilloteaux et al., 2017) in Icy (de Chaumont et al., 2012). Segmentation and rendering was performed in Amira (Thermo Scientific).

## Data availability

Data are available from the corresponding author upon request. Constructs and cell lines are available upon request. Materials involving the ANCHOR system are MTA-restricted and commercially available from NeoVirTech (France).

## Acknowledgements

The ANCHOR system is developed by and commercially available from NeoVirTech (France, www.neovirtech.com). We gratefully acknowledge H. Wodrich (University of Bordeaux, France) and F. Gallardo (NeoVirTech) for helpful discussions. We thank David Bejarano (University Hospital Heidelberg, Germany) for providing pVpr-IN_D64N/D116N_.eGFP. EBFP2-LaminB1-10 was a gift from Michael Davidson (Addgene plasmid # 55244), pSNAP-tag (m) Vector from New England Biolabs & Ana Egana (Addgene plasmid # 101135), pmScarlet_C1 from Dorus Gadella (Addgene plasmid # 85042), psPAX2 from Didier Trono (Addgene plasmid #12260) and pCMV-VSV-G was from Bob Weinberg (Addgene plasmid #8454). We thank Anke-Mareil Heuser and Vera Sonntag-Buck for excellent technical assistance. We would like to acknowledge the microscopy support from the Infectious Diseases Imaging Platform (IDIP) of the Center for Integrative Infectious Disease Research, Heidelberg.

This work was funded by the Deutsche Forschungsgemeinschaft (DFG, German Research Foundation) - Projektnummer 240245660 - SFB 1129 project 5 (H-G.K.), project 6 (B.M.), project 20 (M.L.) and by the TTU HIV in the DZIF (V.L., M.L., H-G.K.).

## Supplementary Figures

**Figure 1 - figure supplement 1.**
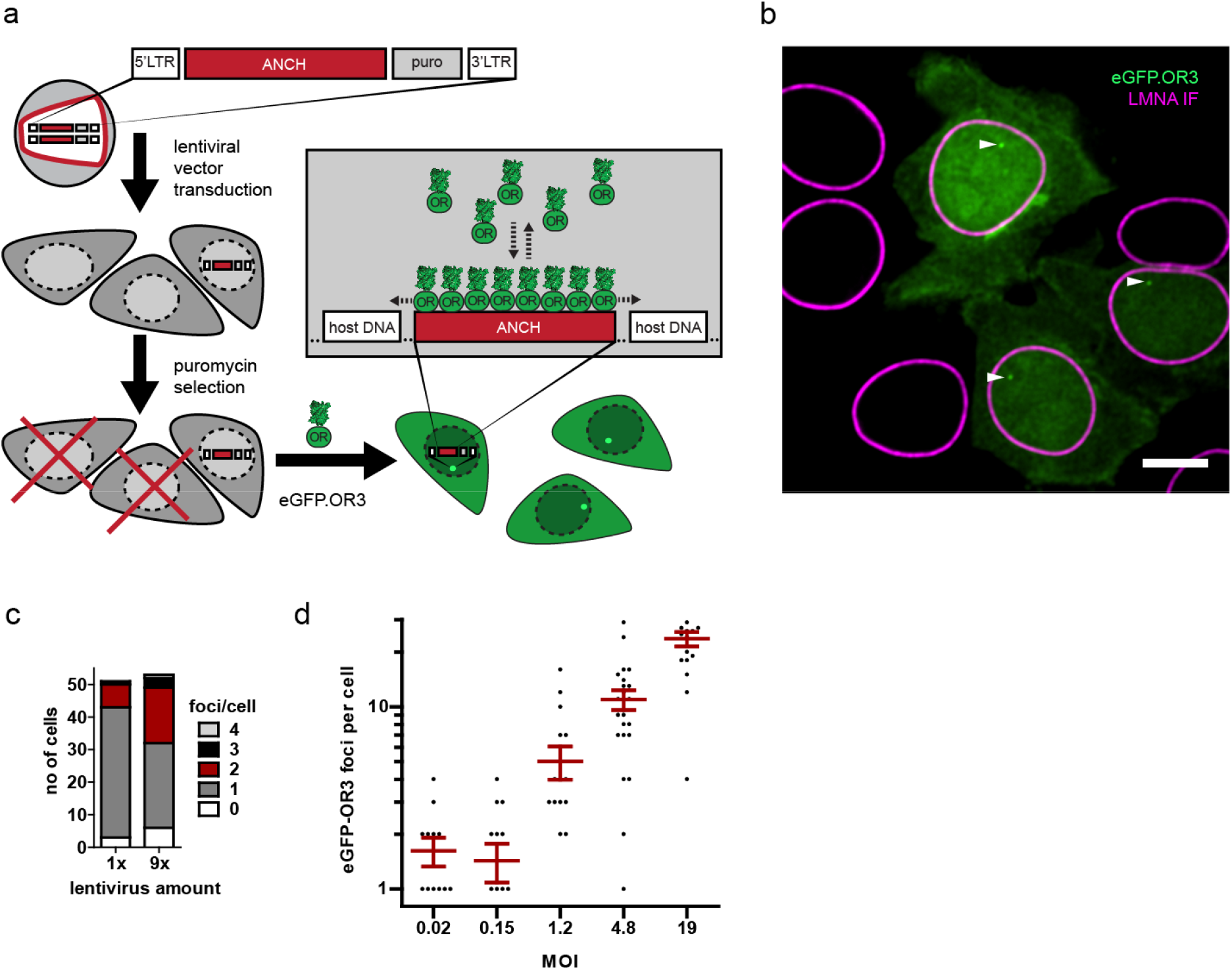
The ANCHOR system enables sensitive visualization of integrated lentiviral DNA without loss through recombination during reverse transcription. (**a**) Scheme of experiment. A HIV-1 lentiviral vector harboring the ANCH sequence and a puromycin expression cassette was used to transduce TZM-bl cells. After 2-3 days puromycin was added to deplete untransduced cells, after which eGFP.OR3 was delivered by lentiviral transduction or transfection. All surviving cells are expected to display one or more OR3 punctae in the nucleus. (**b**) Exemplary cells from low MOI transduction of cells using lentiviral particle containing supernatant. Scale bar: 5 μm. One of two independent experiments is shown. (**c**) Quantification of spots per cell from 50 μl (1×) and 450 μl (9×) used supernatant. (**d**) Titration of concentrated lentiviral particles. MOI was determined by counting the number of surviving cell colonies 5 days after puromycin selection. The transduced cells were kept in culture for over 4 weeks to ensure degradation of unintegrated vDNA products. Error bars represent SEM.

**Figure 1 - figure supplement 2.**
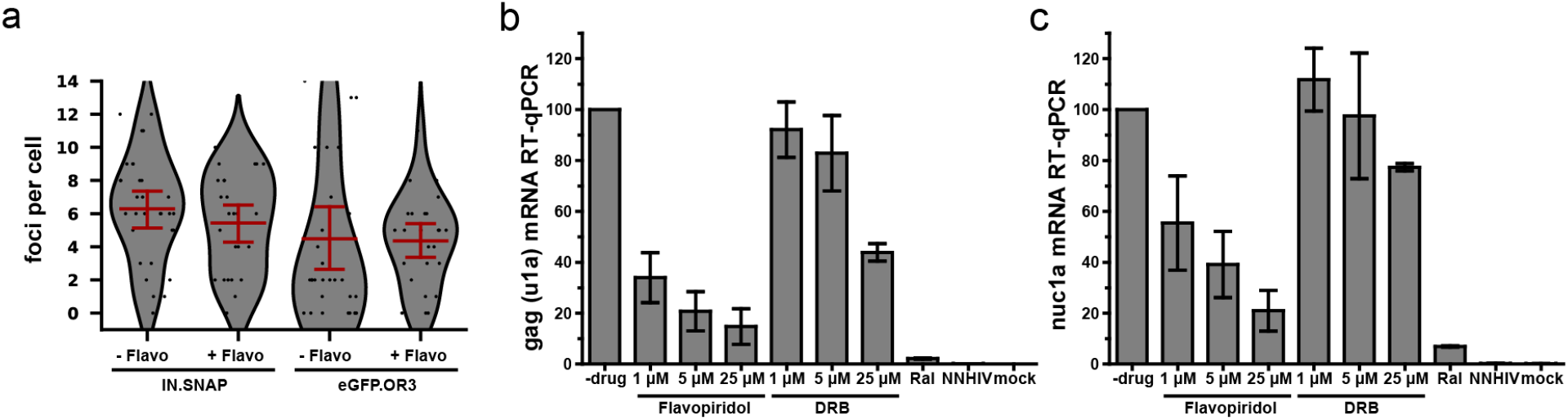
eGFP.OR3 punctae in infected cell nuclei are detected independent of HIV-1 transcription. (**a**) eGFP.OR3 and eBFP2.LMNB1 expressing TZM-bl cells were infected with 30 μU RT/cell VSV-G pseudotyped HIV ANCH for 47 h, after which 5 μM of the pTEF-b transcription initiation inhibitor Flavopiridol was added for another 8 h. Number of nuclear IN.SNAP and eGFP.OR3 punctae were quantified. One of three independent experiments is shown with n > 25 cells per sample, error bars represent 95 % CI. (**b,c**) qRT-PCR of viral RNA products specific for *gag* (u1a) (**b**) and the transcription of the first nucleosome nuc1a (**c**) quantified at 55 h p.i‥ eBFP2.LMNB1 and eGFP.OR3 expressing TZM-bl cells were infected with 5 μU RT/cell VSV-G pseudotyped HIV ANCH. Flavopiridol and the transcription elongation inhibitor 5,6-dichloro-1-beta-D-ribofuranosylbenzimidazole (DRB) were added 8 h prior RNA extraction. The experiment was performed in biological triplicates and error bars represent SD.

**Figure 1 - figure supplement 3.**
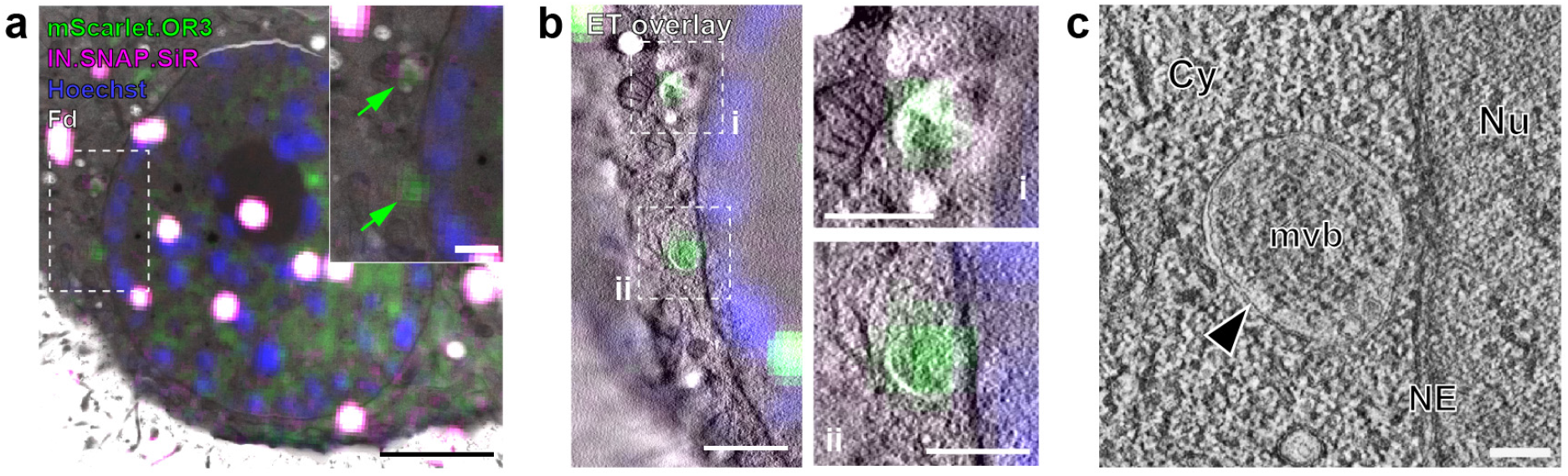
CLEM analysis of infection-independent cytoplasmic OR3 fusion protein accumulations. Experimental details as in Figure 5. (**a**) CLEM overlays (with enlargements) of EM sections of infection-independent mScarlet.OR3 (green) accumulations. Shown is the same cell section as in Figure 5a. (**b**) CLEM-ET overlay of regions enlarged in (**a**). (**c**) Computational slices from tomographic reconstructions at the correlated positions boxed in (**b**, ii). Scale bars: 2.5 μm (**a**), 750 nm (**b**), 500 nm for enlargements (**a,b**) and 100 nm (**c**).

**Figure 3 - figure supplement 1.**
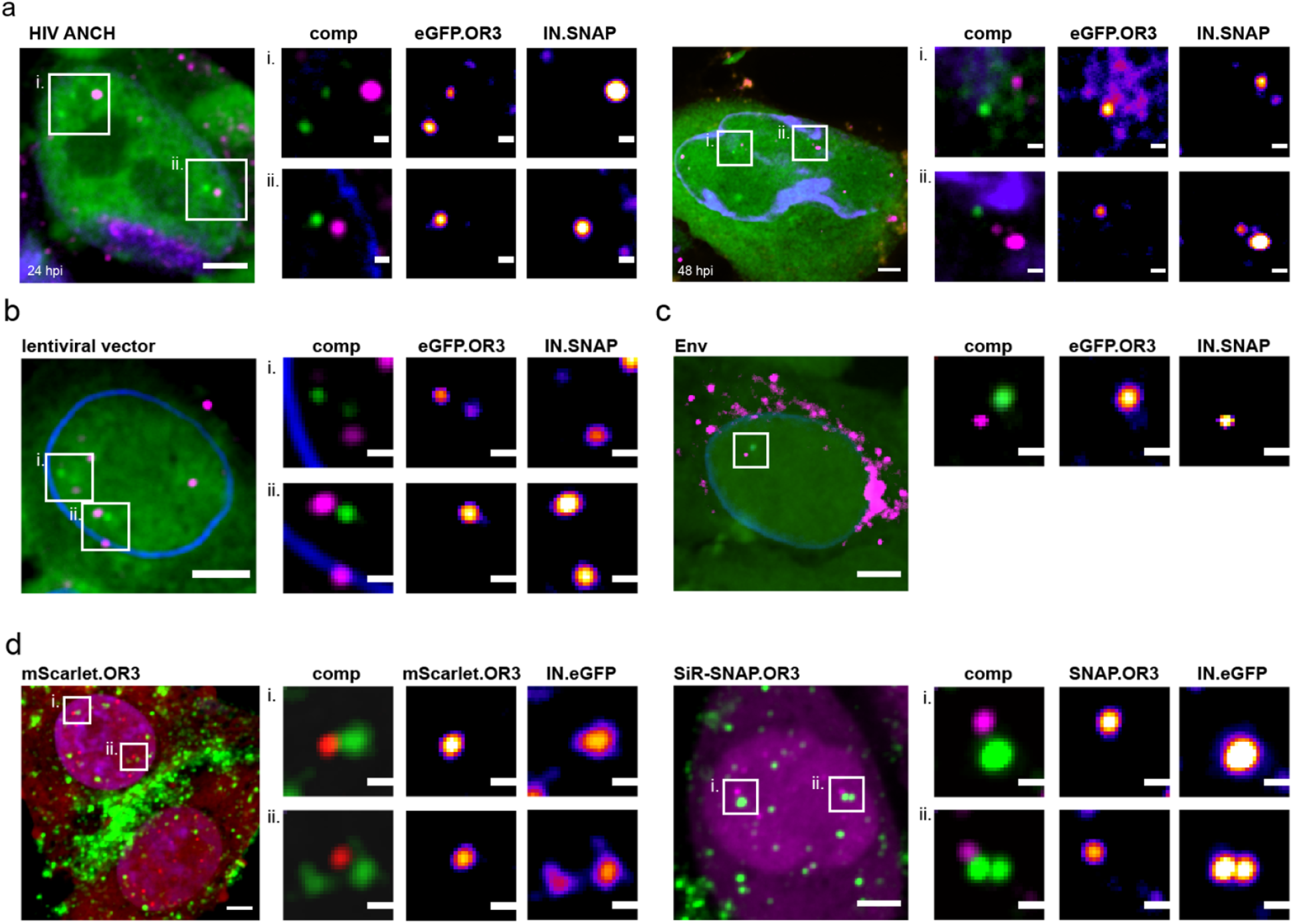
Different TZM-bl cell lines infected with different particles. Infection was performed using 30 μU RT/cell of the following viral particles for 24 h p.i.: Integration competent HIV-1_NL4-3_ ANCH (**a**), an integration competent lentiviral vector (**b**), NNHIV ANCH (**c,d**) pseudotyped with HIV-1 Env fusion protein instead of VSV-G (**c**) or with an inversion of the fluorescent proteins used (**d**). Separation between IN-FP and eGFP.OR3 was observed in all cases. In enlargements OR3 background has been subtracted for clarity. Scale bars: 5 μm (overview) and 1 μm (enlargements); single z plane (**a**), 2.5 μm z-projection (**a-c, d** right panel), z-projection of entire nucleus (**d**left panel).

**Figure 6 - figure supplement 1.**
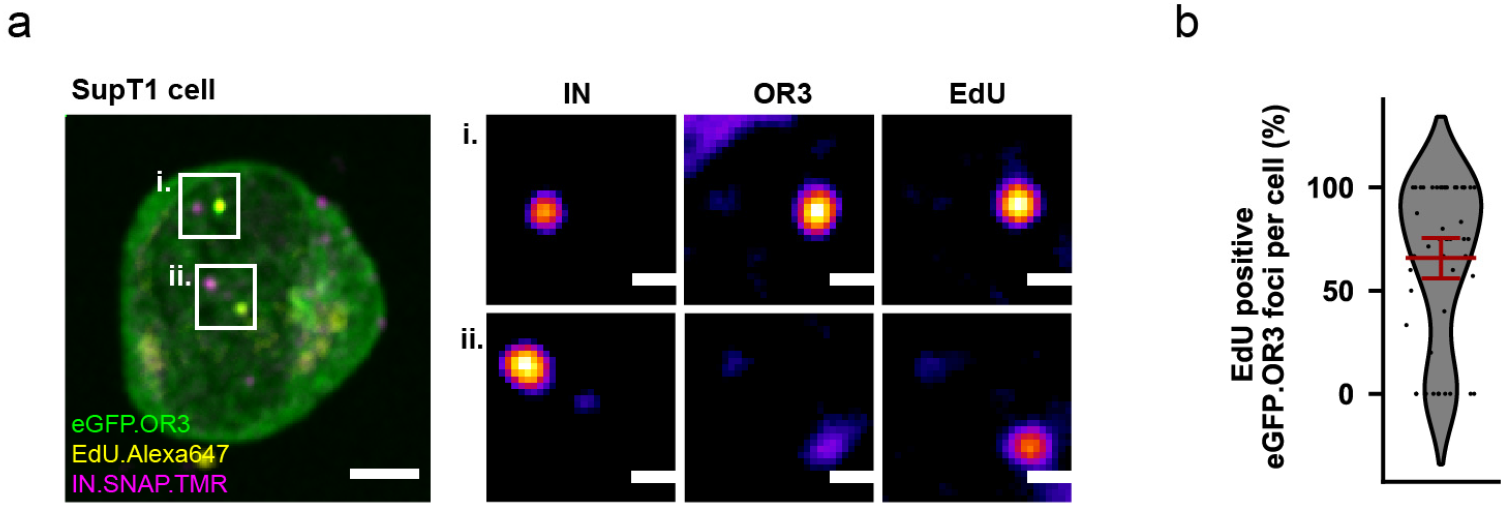
ANCHOR system adopted to the T cell line SupT1. (**a**) SupT1 cells expressing eGFP.OR3 were infected with 30 μU RT/cell VSV-G pseudotyped NNHIV ANCH in the presence of APC and EdU, fixed and click labelled at 24 h p.i‥ A similar separation between IN-FP and eGFP.OR3 as in TZM-bl cells was observed. In enlargements the nuclear background of eGFP.OR3 has been subtracted for clarity. Scale bars: 5 μm (overview) and 1 μm (enlargement). (**b**) Quantification of EdU positive nuclear eGFP.OR3 spots per cell. Pooled data from biological triplicates are shown, error bar represents 95 % CI.

**Video 1: Live cell imaging of viral DNA punctae formation in TZM-bl cells.** Related to Figure 1g and h. While the figure shows raw data (only mean filtered), for the movie 3D stacks were denoised using CARE and rendered in napari (see material and methods for details). The movie is a composite of eBFP2.LMNB1 (magenta) and eGFP.OR3 (green). Scale bar: 5 μm and time in hours:minutes post infection.

**Video 2: Live cell imaging of eGFP.OR3 separation from IN.SNAP.** Related to Figure 3d and e. Shown is a maximum intensity projection of a cell nucleus. The movie is a composite of eBFP2.LMNB1 (blue), eGFP.OR3 (green) and IN.SNAP.SiR (magenta). Scale bar: 5 μm and time in hours:minutes post infection.

**Video 3: Live cell imaging of eGFP.OR3 separation from IN.SNAP.** Related to Figure 3d and e appearance of two eGFP.OR3 punctae from a single IN-FP spot. Shown is a maximum intensity projection of a nuclear particle with the nucleus at top left and the cytoplasm at the bottom right of the movie. The movie is a composite of eBFP2.LMNB1 (blue), eGFP.OR3 (green) and IN.SNAP.SiR (magenta). Scale bar: 2 μm and time in hours:minutes post infection.

**Video 4: CLEM-ET analysis, segmentation and rendering of nuclear capsid cone clustering in HeLa derived cells.** Related to Figure 4e. Morphology of clustered capsid-related structures observed at the position indicated by an IN.SNAP.SiR signal lacking mScarlet.OR3 signal.

**Video 5: CLEM-ET analysis, segmentation and rendering of IN and OR3-positive structures in infected HeLa derived cells.** Related to Figure 5i. Morphology of clustered capsid-related structures observed at the position indicated by co-localizing mScarlet.OR3 and IN.SNAP.SiR.

